# Inducible FAK Deletion but not FAK Inhibition in Endothelial Cells Activates p53 to Suppress Tumor Growth in PYK2-null Mice

**DOI:** 10.1101/2024.11.04.622008

**Authors:** Xiao Lei Chen, Marjaana Ojalill, Christine Jean, Isabelle Tancioni, Shulin Jiang, Antonia Boyer, Duygu Ozmadenci, Sean Uryu, David Tarin, Joseph Schlessinger, Dwayne G. Stupack, David D. Schlaepfer

## Abstract

Focal adhesion kinase (FAK) functions as a signaling and scaffolding protein within endothelial cells (ECs) impacting blood vessel function and tumor growth. Interpretations of EC FAK-null phenotypes are complicated by related PYK2 (protein tyrosine kinase 2) expression, and to test this, we created PYK2^-/-^ FAK^fl/fl^ mice with tamoxifen-inducible EC-specific Cre recombinase expression. At 11 weeks of age, EC FAK inactivation resulted in increased heart and lung mass and vascular leakage only on a PYK2^-/-^ background. Surprisingly, ∼90% of PYK2^-/-^ EC FAK^-/-^ mice survived to 75 weeks of age. Syngeneic melanoma, breast, or lung carcinoma tumors did not grow in PYK2^-/-^ EC FAK^-/-^ mice, but tumors grew normally in PYK2^-/-^ EC FAK^fl/fl^ mice lacking Cre. This tumor inhibitory phenotype was associated with abortive EC vessel sprouting, enhanced EC p53 tumor suppressor and p21CIP1 (cyclin-dependent inhibitor 1) expression, and alterations in serum cytokine levels. To discern the role of FAK kinase versus scaffolding activity in ECs, we generated kinase defective (FAK K454R, KD) PYK2^-/-^ EC FAK^fl/KD^ and PYK2^-/-^ EC FAK^fl/WT^ (WT, wildtype) mice. Hemizygous EC FAK^-/KD^ expression supported primary tumor growth but not metastasis, implicating EC FAK activity in tumor dissemination. *In vitro*, hemizygous expression of either WT or KD FAK suppressed EC p21CIP1 levels and cell death observed in primary PYK2^-/-^ EC FAK^-/-^ ECs. Combined FAK and PYK2 knockdown in tumor cells also increased p21CIP1 and PARP1 (poly ADP-ribose polymerase 1) levels in a p53-associated manner impacting anchorage-independent growth. Together, these results underscore the linkage between PYK2 and FAK loss with p53 activation impacting tumor growth.

**Impact Statement:** PYK2-null combined with endothelial cell-specific FAK transgenic mouse models show that loss of FAK activity limits tumor spread and that genetic or chemical degradation preventing combined FAK-PYK2 expression may be an approach to induce a p53-associated anti-tumor response.

## Introduction

Focal adhesion kinase (FAK, *PTK2*) and protein tyrosine kinase 2 beta (PYK2, *PTK2B*) are evolutionarily related non-receptor protein-tyrosine kinases^1,2^. FAK is ubiquitously expressed and best known to facilitate cell surface integrin receptor signaling promoting cell migration, mechano-sensing, and survival^3,4^. PYK2 also exhibits a relatively broad expression, with key roles in neuronal and myeloid cell signaling^5^. Both PYK2 and FAK are activated in response to a broad range of stimuli, and agents which inhibit these kinases are being evaluated in clinical trials^6–8^.

Angiogenic endothelial cells (ECs) form blood vessel conduits^9,10^ and both FAK and PYK2 are co-expressed in murine and human ECs^11,12^. Notably, EC FAK expression and intrinsic kinase activity are essential for developmental vasculogenesis^13–15^ and pharmacological or genetic FAK inhibition can limit tumor angiogenesis in mice^16–20^. In ECs, vascular endothelial growth factor facilitates FAK activation and recruitment to cell-cell adherens junctions, binding to VE-cadherin, and β-catenin tyrosine phosphorylation within a VE-cadherin complex in the control of vascular permeability and tumor metastasis^21,22^. Interestingly, PYK2 is activated and recruited to cell-cell junctions following the disruption of VE-cadherin homotypic binding and supports pulmonary angiogenesis^23^. Additionally, the binding and phosphorylation of p130Cas by FAK or PYK2 functions as a parallel signaling pathway supporting angiogenesis^24^.

Murine FAK knockout or genetic inactivation of FAK activity results in developmental lethality^25^. In contrast, PYK2 knockout mice are viable and fertile, but display altered lymphoid, myeloid and macrophage signaling responses^25,26^. PYK2 possesses a similar domain structure and regulatory phosphorylation sites as FAK^8^. Further, PYK2 binds many of the same adaptor proteins as FAK, yet endogenous or exogenous PYK2 expression does not rescue motility defects of FAK-null primary fibroblasts^27^. Importantly, an engineered, focal adhesion-targeted PYK2 chimeric protein rescued FAK-null cell migration equally as FAK re-expression^28^, suggesting a key difference may be in the intracellular targeting of these related proteins.

However, one conserved “signaling” role for FAK and PYK2 is to monitor and limit p53 tumor suppressor levels in response to cellular stress^29^. During development, FAK inactivation leads to elevated p53-mediated transcription of p21CIP1 (cyclin-dependent kinase inhibitor 1) and prevention of primary cell growth^30^. This p53- and p21-dependent growth arrest is controlled in part by FAK and/or PYK2-mediated nuclear localization, p53 binding, and enhanced p53 turnover by proteasomal degradation^31^ in a manner independent of intrinsic FAK-PYK2 kinase activity. In addition to canonical cell intrinsic p53-mediated effects in promoting cell cycle arrest, apoptosis, or cell senescence as barriers for cancer development^32,33^, p53 activation can also suppress tumorigenesis in a cell extrinsic manner by promoting an anti-tumor microenvironment^34,35^. However, it remains unclear whether FAK and PYK2 function as redundant modulators of p53 activity beyond development.

To bypass developmental restriction points requiring FAK, several independent Cre-recombinase models of inducible floxed (fl) FAK inactivation in adult mice have been created^36–38^. Surprisingly, conditional FAK loss within ECs in one mouse model still supported normal angiogenesis^39^ whereas in another EC knockout model, FAK loss inhibited tumor neovascularization^17^. As ECs possess the adaptive capacity to switch to PYK2-dependent survival signaling upon genetic FAK inactivation^31^, and PYK2 expression was detected in FAK-null ECs^39^, it is conceivable that FAK and PYK2 may share overlapping functions within ECs in supporting p53 regulation, tumor growth and angiogenesis.

To test this hypothesis, we made PYK2^-/-^ FAK^fl/fl^ mice and back-crossed to C57Bl6 strain purity. Using a tamoxifen-inducible EC-specific Cre, FAK inactivation on a PYK2^-/-^ background resulted in increased heart and lung tissue wet weight with vascular leakage but limited lethality of tamoxifen treated Cre^+^ (FAK loss) but not Cre^-^ littermate mice. Loss of EC FAK in PYK2^-/-^ mice prevented syngeneic breast, melanoma, and lung carcinoma tumor growth.

Mechanistically, tumor inhibition was associated with EC-intrinsic elevation of p53, p21CIP1 and markers of DNA damage as well as EC-extrinsic effects altering peripheral blood cytokine levels. Inhibition of EC FAK activity in PYK2^-/-^ mice did not activate p53 or prevent tumor growth, but chemical degradation of FAK and PYK2 protein activated p53 signaling also in tumor cells. Together, our combined results define distinct phenotypic differences between the loss of FAK-Pyk2 expression compared to the inhibition of FAK-Pyk2 activity.

## Results

### Inactivation of FAK within adult mouse blood vessel ECs of PYK2^-/-^ mice

EC inactivation of FAK expression or activity results in embryonic developmental lethality associated with vascular defects^13,14^. Lethality is averted upon FAK inactivation after 6 weeks of age and in these mice, EC FAK function is implicated in promoting angiogenesis^17^. However, PYK2 expression is elevated or maintained upon FAK knockdown in multiple cell types *in vitro* and in different conditional FAK knockout mouse models *in vivo*^31,39,40^. Double or single knockout of PYK2 and FAK within intestinal epithelial cells revealed essential functions in barrier maintenance and epithelial repair following injury^41^. In ECs, as PYK2 can promote sprouting under conditions of limited FAK activation^42^, we performed crosses with PYK2^-/-^ and FAK^fl/fl^ mice containing a tamoxifen-activatable stem cell leukemia (SCL) promoter-driven EC-specific Cre recombinase (Cre-ERT) to create a double knockout cell model *in vivo* (Figure 1). In functional studies, Cre^-^ littermates were used as controls.

**Figure 1.**
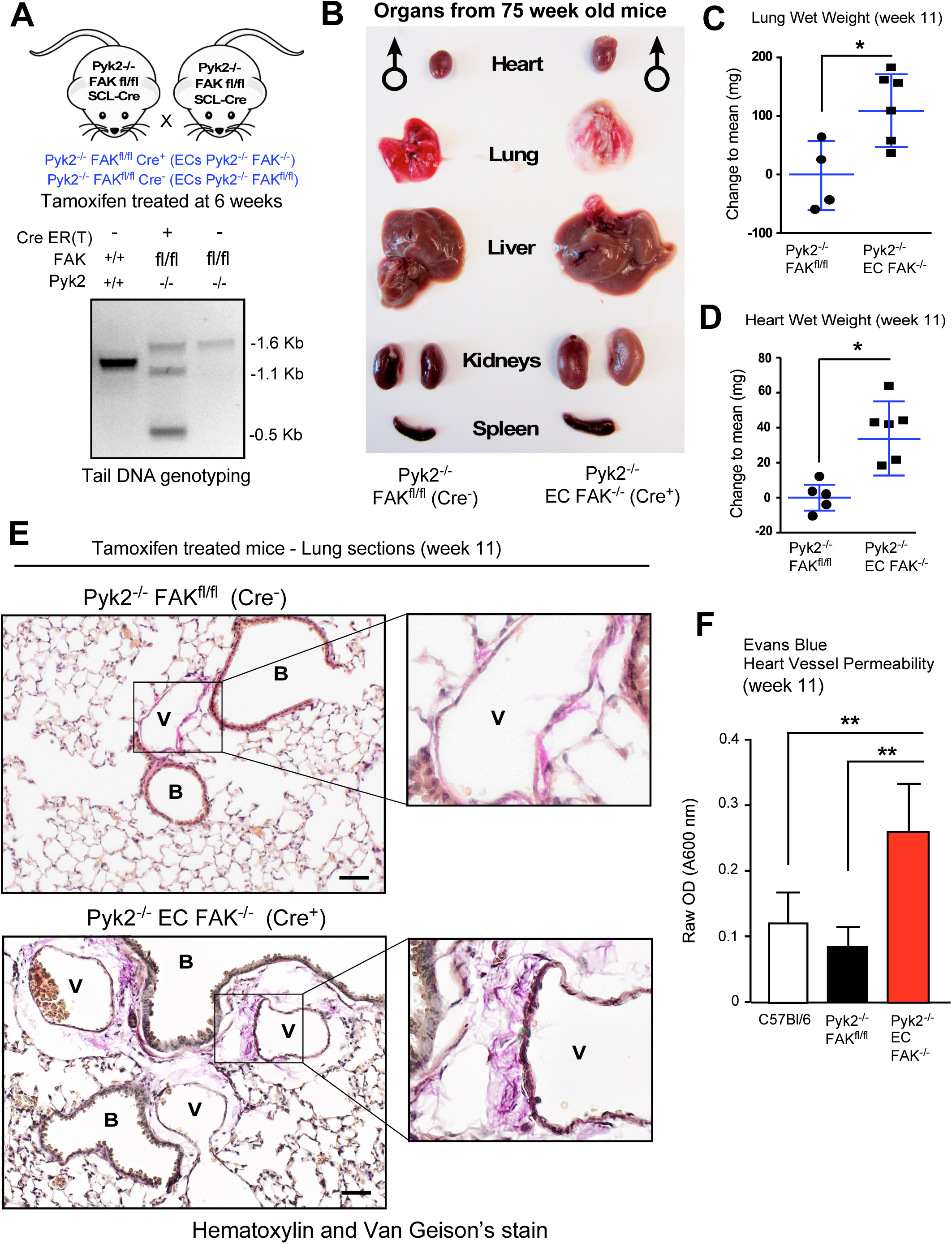
Vascular-associated tissue edema and fibrosis in mice upon inducible EC FAK loss on a PYK2^-/-^ background. (**A**) Schematic of mouse crosses. The stem cell leukemia (SCL) promoter-driven Cre recombinase fusion with the estrogen receptor allows for tamoxifen-induced Cre activation (ERT) in ECs. As SCL-Cre-ERT is a single allele, littermate PYK2^-/-^ FAK^fl/fl^ offspring will be either Cre^+^ or Cre^-^. Below, representative tail DNA genotyping for FAK by PCR, 1.5 kB FAK^+/+^, 1.6 kb FAK^fl/fl^, and 1.1 plus 0.5 kB for FAK^fl/fl^ with Cre. (**B**) At 6 weeks, mice were given tamoxifen chow for 3 weeks, regular chow for 1 week, and then tissues analyzed at times indicated. Representative images of non-perfused tissues from male PYK2^-/-^ FAK^fl/fl^ Cre^-^ and PYK2^-/-^ EC FAK^-/-^ littermate mice at 75 weeks of age. (**C**) Mean lung and (**D**) heart wet weight from tamoxifen-treated PYK2^-/-^ FAK^fl/fl^ Cre^-^ and PYK2^-/-^ EC FAK^-/-^ littermate mice at ∼11 weeks of age. Points are from individual mice, mean weight of C57Bl6 tissues was set to 0 (+/- SD, * *P* <0.05, T-test). (**E**) Representative hematoxylin plus van Gieson (collagen fiber, pink) staining of perfused, formalin fixed and paraffin-embedded lung sections from tamoxifen-treated mice with the indicated genotypes at ∼11 weeks of age. Inset, enlargement of selected vessel (B, bronchiole and V, vessels). Some vessels contain red blood cells due to incomplete saline perfusion. Scale is 25 µm. (**F**) Basal heart tissue vascular permeability was measured in tamoxifen-treated mice of the indicated genotype. Tissue accumulation of circulating Evans blue dye was measured 30 min after tail-vein injection. Shown are mean values from two independent experiments (+/- SD, n=6, ** *P* <0.01) by ANOVA with Tukey’s post-hoc test.

Littermates with PYK2^-/-^ FAK^fl/fl^ SCL Cre^+^ or Cre^-^ genotypes were provided tamoxifen chow at week 6 for 3 weeks, returned to normal chow for 1 week, and then evaluated at week 11. Tail genotyping confirmed FAK^fl/fl^ allele inactivation *in vivo* (Figure 1A). Heart and lung tissue lysates, which are greatly enriched for ECs, showed reduced total FAK protein levels in PYK2^-/-^ EC FAK^-/-^ mice (Figure 1 - figure supplement 1A and 1B). Confirmation of EC-FAK knockout was performed by co-staining for CD31 (EC marker) and FAK within heart tissue by immunofluorescence (Figure 1 - figure supplement 1C). These results are consistent with EC FAK being at least initially compatible with adult mice lacking PYK2, though approximately 10% of PYK2^-/-^ EC FAK^-/-^ mice died by week 11.

Despite an absence of visible phenotypes observed in PYK2^-/-^ EC FAK^-/-^ mice at week 11, these mice exhibited enlarged lungs, hearts, livers, and kidneys compared to Cre^-^ PYK2^-/-^ FAK^fl/fl^ littermates, which was detected upon macroscopic tissue examination (Table 1). While this phenotype was associated with increased early mortality, some PYK2^-/-^ EC FAK^-/-^ survived to old age (75 weeks) with enlarged organs and increased visceral adipose tissue compared to Cre^-^ littermates (Figure 1B and Figure 1 - figure supplement 1D). Consistent with the enlarged tissue phenotype, dissected lung and heart wet weight was elevated significantly in PYK2^-/-^ EC FAK^-/-^ mice compared to Cre^-^ littermates at week 11 (Figures 1C and D). Immunohistochemical staining of lungs revealed enlarged “spaces” and elevated collagen deposition surrounding PYK2^-/-^ EC FAK^-/-^ vessels, possibly associated with tissue edema as compared to other genotypes (Figure 1E and Figure 1 – figure supplement 2). Elevated lung collagen staining was observed in lungs of PYK2^-/-^ EC FAK^-/-^ compared to C57BL/6 mice (Figure 1 – figure supplement 2) and increased vascular space was also observed in hearts of PYK2^-/-^ EC FAK^-/-^ mice at week 11 compared to controls (Figure 1 – figure supplement 3). To determine if increased tissue mass was associated with vascular leakage, basal vascular heart permeability differences were quantified by tissue accumulation of Evans blue dye after tail vein administration (Figure 1F).

**Table 1.** Phenotypes observed after tamoxifen treatment of FAK^fl/fl^ SCL-Cre^-^ and SCL-Cre^+^ mice on a Pyk2^-/-^ C57BL/6 background at 11 weeks of age.

| Phenotype | FAK<sup>fl/fl</sup> | EC FAK<sup>-/-</sup> |
| --- | --- | --- |
| Enlarged Lungs | 1/10 | 11/16 |
| Enlarged Heart | 0/10 | 4/16 |
| Enlarged Liver | 1/10 | 8/16 |
| Enlarged Kidneys | 0/10 | 5/16 |
| Death | 0/10 | 2/18 |

Significantly elevated basal vascular leakage occurred in PYK2^-/-^ EC FAK^-/-^ mice compared to controls. Taken together, our results support the notion that PYK2 and FAK loss within ECs weakens blood vessel barrier integrity, but this does not necessarily lead to premature lethality.

### Inducible loss of EC FAK in PYK2^-/-^ mice limits implantable melanoma tumor formation

We previously established a compensatory role for PYK2 in supporting angiogenesis upon EC FAK loss *in vivo*^39^. To determine the effects of combined PYK2 and FAK loss within adult mouse vessel ECs on tumor growth, red fluorescent protein (RFP) expressing B16F10 melanoma cells were injected subcutaneously into PYK2^-/-^ EC FAK^-/-^ and Cre^-^ littermate control mice, and after 5 days, all mice were given tamoxifen chow and monitored for changes in tumor size (Figure 2A). Tumor volume measurements trended lower in PYK2^-/-^ EC FAK^-/-^ mice (Figure 2B) and after 21 days, final B16F10 tumor mass was significantly less when grown in PYK2^-/-^ EC FAK^-/-^ compared to Cre^-^ control mice (Figure 2C).

**Figure 2.**
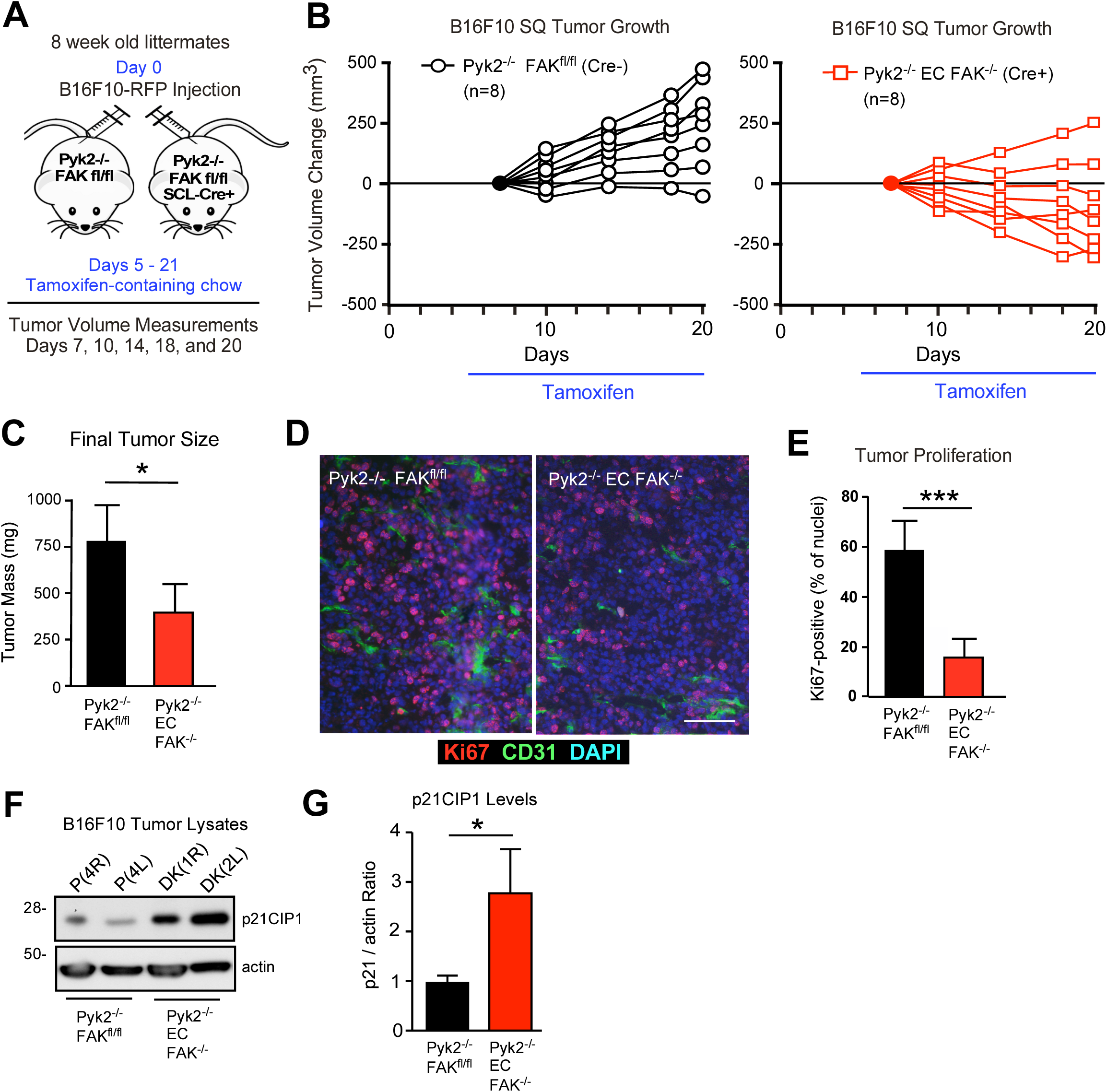
Inducible EC FAK loss on a PYK2^-/-^ background slows syngeneic B16F10 melanoma tumor growth. (**A**) Schematic of B16F10 tumor cell injection at Day 0, change to tamoxifen chow at Day 5, and caliper tumor measurements at indicated days. (**B**) Mean tumor volume normalized at Day 7 as shown for individual mice (n=8 mice per group) for PYK2^-/-^ FAK^fl/fl^ Cre^-^ (black) and PYK2^-/-^ EC FAK^-/-^ Cre^+^ (red) through Day 20. Results are from one of two independent experiments. (**C**) Final mean tumor mass at Day 21 (+/- SD, *P* <0.05 T-Test). (**D**) Representative immunofluorescence staining of B16F10 tumor sections from PYK2^-/-^ FAK^fl/fl^ Cre^-^ and PYK2^-/-^ EC FAK^-/-^ mice with antibodies to Ki67 (red), CD31(green), and DAPI (blue). Scale is 25 µm. (**E**) Quantitation of CD31 vessel EC staining. Mean percentage of CD31 and DAPl-positive cells per field (+/- SD, 10 sections from 3 tumors for each group, *** *P* <0.001 T-Test). (**F**) Tumor protein lysates collected at Day 20 were immunoblotted for p21CIP1 and actin. Lanes represent B16F10 tumor lysates from individual mice of the indicated genotype. (**G**) Quantitation of p21CIP1 to actin protein levels in B16F10 tumor lysates. Values (n=4 each) means (+/- SD, * *P* <0.05, T-Test).

As a measure of cell proliferation, Ki67 immunohistochemical staining was significantly decreased in B16F10 tumors grown in tamoxifen treated PYK2^-/-^ EC FAK^-/-^ compared to PYK2^-/-^ FAK^fl/fl^ Cre^-^ control mice (Figures 2D and E). Ki67-positive tumor cells were primarily detected nearby CD31-stained ECs at the tumor periphery in PYK2^-/-^ EC FAK^-/-^ mice whereas Ki67 staining was detected throughout B16F10 tumors grown in PYK2^-/-^ FAK^fl/fl^ Cre^-^ mice (Figure 2D). Surprisingly, p21CIP1 protein levels were increased compared to actin in lysates of the smaller B16F10 tumors from PYK2^-/-^ EC FAK^-/-^ compared to larger B16F10 tumors from PYK2^-/-^ FAK^fl/fl^ mice (Figures 2F and G). Although we did not perform sequence verification, B16F10 cells express a mutant form of p53, and increased p21CIP1 cell cycle inhibitor expression is regulated in part by wildtype p53 activity^43^. Thus, increased p21CIP1 levels may not be a tumor intrinsic response. Nonetheless, our results support the hypothesis that induced loss of EC FAK on a PYK2^-/-^ background is associated with EC cell-intrinsic or -extrinsic changes (or both) and the possible switch from a permissive to repressive host tumor microenvironment.

### PYK2- and FAK-null ECs express elevated p53 and p21CIP1 levels *in vivo*

To determine if markers of p53 activation occur in lung or heart vessels of non-tumor bearing PYK2^-/-^ EC FAK^-/-^ mice, protein immunoblotting of tissue lysates and immunohistochemical staining was performed after tamoxifen treatment of Cre^+^ and Cre^-^ PYK2^-/-^ FAK^fl/fl^ mice (Figure 3). Elevated levels of p53, p16INK4A tumor suppressor, p21CIP1 cell cycle inhibitor, and HP1-γ, a chromatin binding protein marker of DNA damage were detected in heart lysates of Cre^+^ PYK2^-/-^ EC FAK^-/-^ compared to Cre^-^ PYK2^-/-^ FAK^fl/fl^ mice. (Figure 3A). Elevated p53 (Figure 3B) and p21CIP1 (Figure 3C) staining was detected in small blood vessels within lungs of PYK2^-/-^ EC FAK^-/-^ compared to Cre^-^ control mice (Figure 3D and E). Co-staining of lung and heart sections for CD31, TUNEL, and DAPI also revealed increased levels of programmed cell death in Cre^+^ PYK2^-/-^ EC FAK^-/-^ relative to Cre^-^ mice (Figure 3 – figure supplement 1). These results support the notion that induced loss of FAK within ECs on a PYK2-null background generates a p53-associated intrinsic cellular stress signal and increased levels of markers associated with DNA damage.

**Figure 3.**
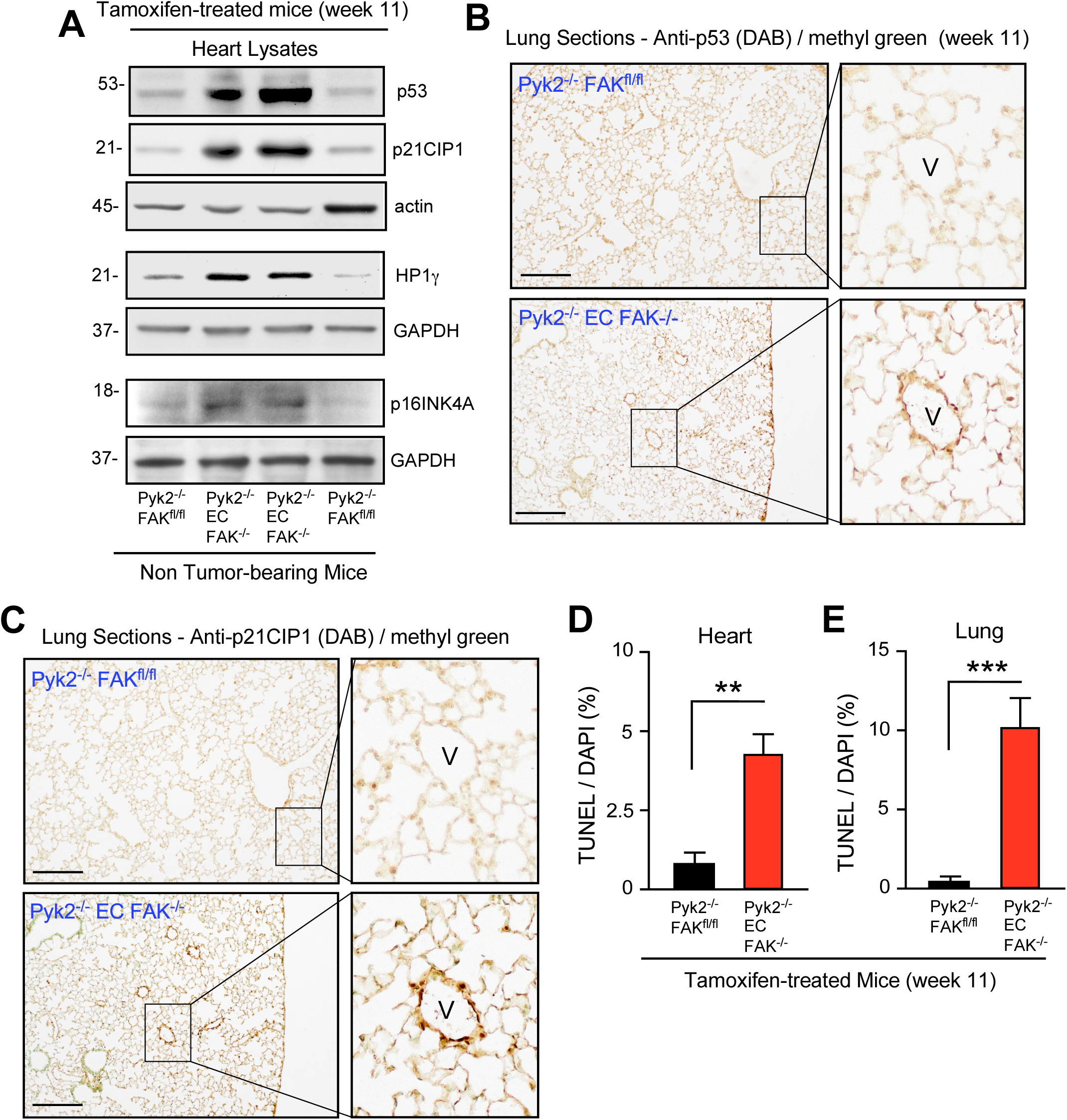
Elevated p53 tumor suppressor and p21CIP1 levels in heart lysates and within lung vessels of non-tumor bearing PYK2^-/-^ EC FAK^-/-^ mice. (**A**) Heart protein lysates from tamoxifen-treated PYK2^-/-^ EC FAK^-/-^ and PYK2^-/-^ FAK^fl/fl^ Cre^-^ mice (n=2) were immunoblotting for p53, p21CIP1, and actin; HP1γ and GAPDH; and p16INK4A and GAPDH. (**B**) and **(C**) Representative immunostaining of paraffin-embedded lung sections for p53 (panel B) or p21CIP1 (panel C) from tamoxifen-treated PYK2^-/-^ EC FAK^-/-^ and PYK2^-/-^ FAK^fl/fl^ Cre-mice. Antibodies were detected by 3,3’-diaminobenzidine (brown) and slides were counter-stained with methyl green. Scale is 200 µm. Inset, lung blood vessel (V). (**D**) and (**E**) Quantitation of TUNEL staining within heart (D) or lung (E) vessels from tamoxifen-treated mice at 11 weeks of age. Mean percentage of DAPl-positive cells per field (+/-SD, 10 sections from each genotype, ** *P* <0.01, *** *P* <0.001 by T-Test).

### Hemizygous EC FAK-KD expression prevents spontaneous vascular permeability on a PYK2^-/-^ background

During development, FAK-KD knock-in mutation results in early embryonic lethality associated with vessel morphogenic defects^14^. Previously, using the SCL-Cre-ERT model and FAK^fl/fl^ mice, we showed that creating mice hemizygous for EC FAK^-/KD^ expression prevented spontaneous B16F10 melanoma metastasis but not tumor growth^22^. To extend these results, Cre^+^ PYK2^-/-^ FAK^fl/fl^ mice were crossed with heterozygous FAK^WT/KD^ mice to yield Pyk2^-/-^ EC FAK^fl/WT^ and Pyk2^-/-^ EC FAK^fl/KD^ littermates (Figure 4A). Tail genotyping was used to verify *in vivo* Cre targeting of FAK (0.5 kb band generation) after tamoxifen treatment (Figure 4B).

**Figure 4.**
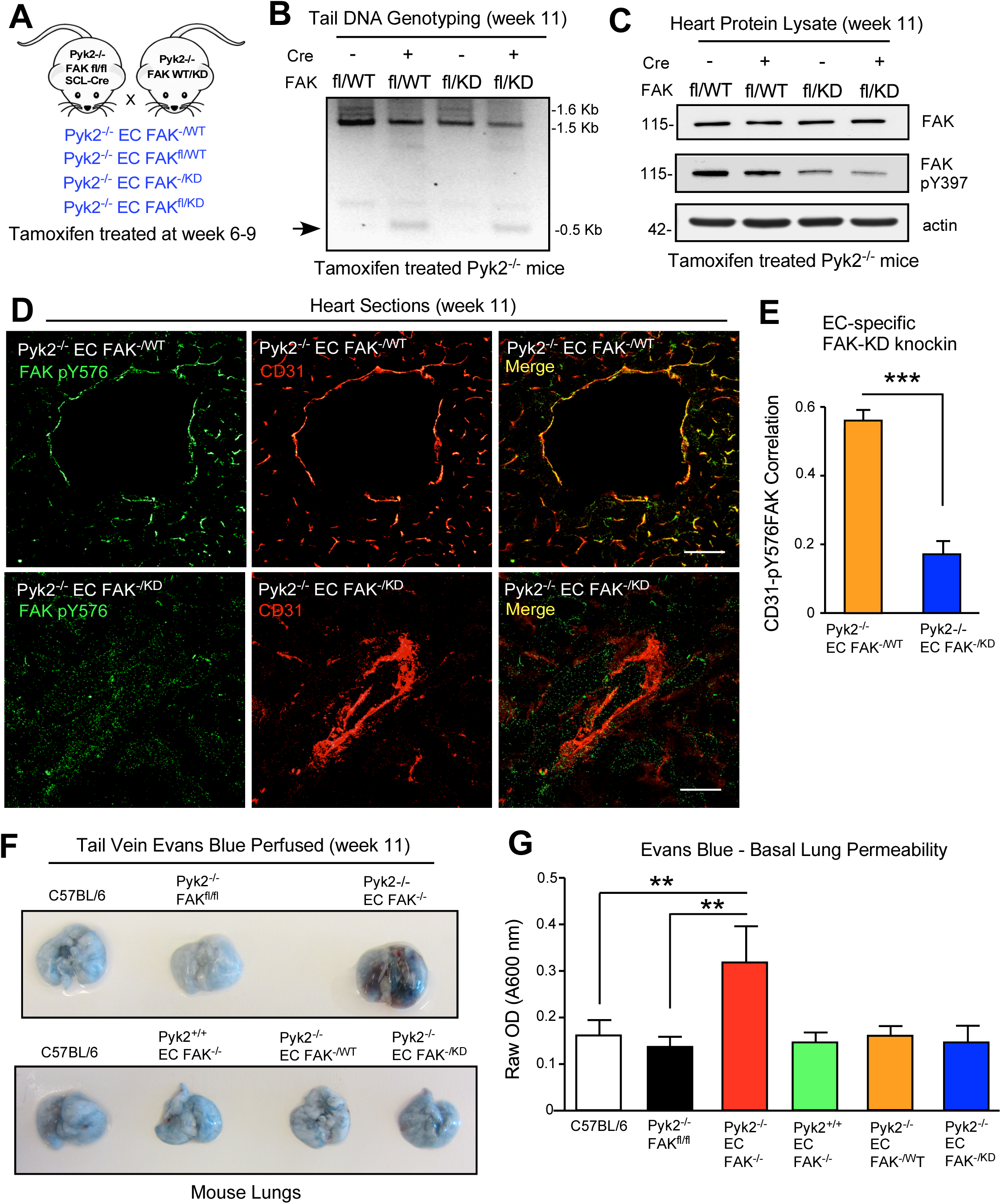
Hemizygous kinase-deficient (KD, K454R) FAK expression in ECs prevents spontaneous vascular permeability on PYK2^-/-^ mouse background. (**A**) Schematic of PYK2^-/-^ FAK^fl/fl^ SCL-Cre crosses with heterozygous FAK^WT/KD^ knockin mice and with tamoxifen treatment of littermates initiated at 6 weeks of age. (**B**) Representative PCR tail DNA genotyping for Cre recombinase targeting the single floxed FAK allele (500 bp) in tamoxifen-treated mice with or without SCL-Cre. (**C**) Representative FAK, pY397 FAK, and actin immunoblotting of heart protein lysates at 11 weeks of age from tamoxifen treated Cre^+^ or Cre^-^ PYK2^-/-^ FAK^fl/WT^ and PYK2^-/-^ FAK^fl/KD^ mice. (**D**) Immunostaining of frozen heart sections for ECs (CD31, red) and activated FAK (pY576 FAK, green) from tamoxifen-treated PYK2^-/-^ EC FAK^-/WT^ and PYK2^-/-^ EC FAK^-/KD^ mice. Merged images show FAK pY576 staining in association with ECs (yellow) only in hearts of PYK2^-/-^ EC FAK^-/WT^ mice. Scale is 25 μm. (**E**) Cell Profiler correlation value of CD31 and pY576 FAK staining overlap within heart sections shown in panel D. Values are means from 10 images, n= 2 mice each genotype, +/- SD *** p<0.01, T-Test). Complete overlap = 1.0. (**F**) Representative images of Evan’s blue dye leakage in lungs of tamoxifen-treated mice of the indicated genotype at 3 h after tail vein injection. (**G**) Quantification of Evans blue dye in lung tissues. Values are means +/- SD from two independent experiments (n=6-8 mice per group, ** *P* <0.01, ANOVA with Tukey’s post hoc test).

Heart lysates showed reduced FAK Y397 phosphorylation from PYK2^-/-^ EC FAK^-/KD^ compared to PYK2^-/-^ EC FAK^-/WT^ mice (Figure 4C). To validate the reduction in FAK activity in vessel-associated ECs, heart sections were co-stained with antibodies to CD31 and with an antibody reporter of active FAK (phosphorylated at FAK Tyr-576)^44^. Whereas heart ECs of PYK2^-/-^ EC FAK^-/WT^ mice exhibited high levels of CD31 and FAK pY576 co-staining, this was significantly reduced in CD31-positive heart vessels of PYK2^-/-^ EC FAK^-/KD^ mice. Together, these results provide validation for selective genetic FAK inhibition but not alteration of FAK expression in PYK2^-/-^ EC FAK^-/KD^ mice (Figures 4D and E).

Interestingly, heart and lung tissue appeared phenotypically normal in PYK2^-/-^ EC FAK^-/KD^ and PYK2^-/-^ EC FAK^-/KD^ mice despite reduced FAK Y576 phosphorylation (Figure 4, table supplement 1 and figure supplement 1A and B). No detectable immunohistochemical expression changes of p53 or p21CIP1 in vessels were detected in PYK2^-/-^ EC FAK^-/KD^ mice compared to PYK2^-/-^ EC FAK^-/WT^ mice (Figure 4 – figure supplement 1C and D) as occurred in lung vessels of PYK2^-/-^ EC FAK^-/-^ compared to Cre^-^ mice (Figure 3). To determine if single allele (hemizygous) FAK-WT or FAK-KD expression rescued vascular leakage observed in PYK2^-/-^ EC FAK^-/-^ mice (Figure 1), Evans blue dye accumulation in lung tissue was quantified in mice of different genotypes (Figure 4F and G). No differences in basal permeability were observed between PYK2^+/+^ EC FAK^-/-^, PYK2^-/-^ EC FAK^-/WT^, or PYK2^-/-^ EC FAK^-/KD^ and normal C57Bl6 mice. Increased lung vessel permeability in PYK2^-/-^ EC FAK^-/-^ mice was accompanied with significantly increased stromal γ-H2AX pSer139 staining (a marker of DNA damage) surrounding smooth muscle actin-positive lungs vessels compared to lungs from control PYK2^-/-^ FAK^fl/fl^ and PYK2^-/-^ EC FAK^-/KD^ mice (Figure 4 – figure supplement 2). Taken together, these results support the notion that FAK expression, but not intrinsic FAK activity, is both sufficient and required for maintenance of vascular barrier function.

### EC FAK-KD expression supports tumor growth and limits metastasis in PYK2^-/-^ mice

Past transgenic mouse studies found that EC FAK loss did not alter pancreatic ductal carcinoma tumors^45^ and that EC FAK-KD expression did not prevent melanoma tumor growth^19,22^. One interpretation of these results is that EC FAK activity may not be essential for supporting tumor growth. An alternate interpretation is that EC PYK2 expression may have compensatory function in the absence of FAK. To better resolve this, subcutaneous B16F10 tumor growth was evaluated upon inactivation of EC FAK expression or activity on a PYK2^-/-^ background (Figure 5). Tumor growth in PYK2^-/-^ EC FAK^-/KD^ mice over 20 days was varied and was similar to that in PYK2^-/-^ EC FAK^-/WT^ mice in some assays (Figure 5A), but significantly less in repeated studies (Figure 5B). Yet consistently, and strikingly, B16F10 cells remained as very small tumor clumps in PYK2^-/-^ EC FAK^-/-^ mice over 20 days and this was accompanied by the lack of detectable spontaneous metastasis to auxiliary and inguinal lymph nodes as observed in 85% of B16F10 tumor-bearing C57Cl6 control mice (Figure 5 – figure supplement 1).

**Figure 5.**
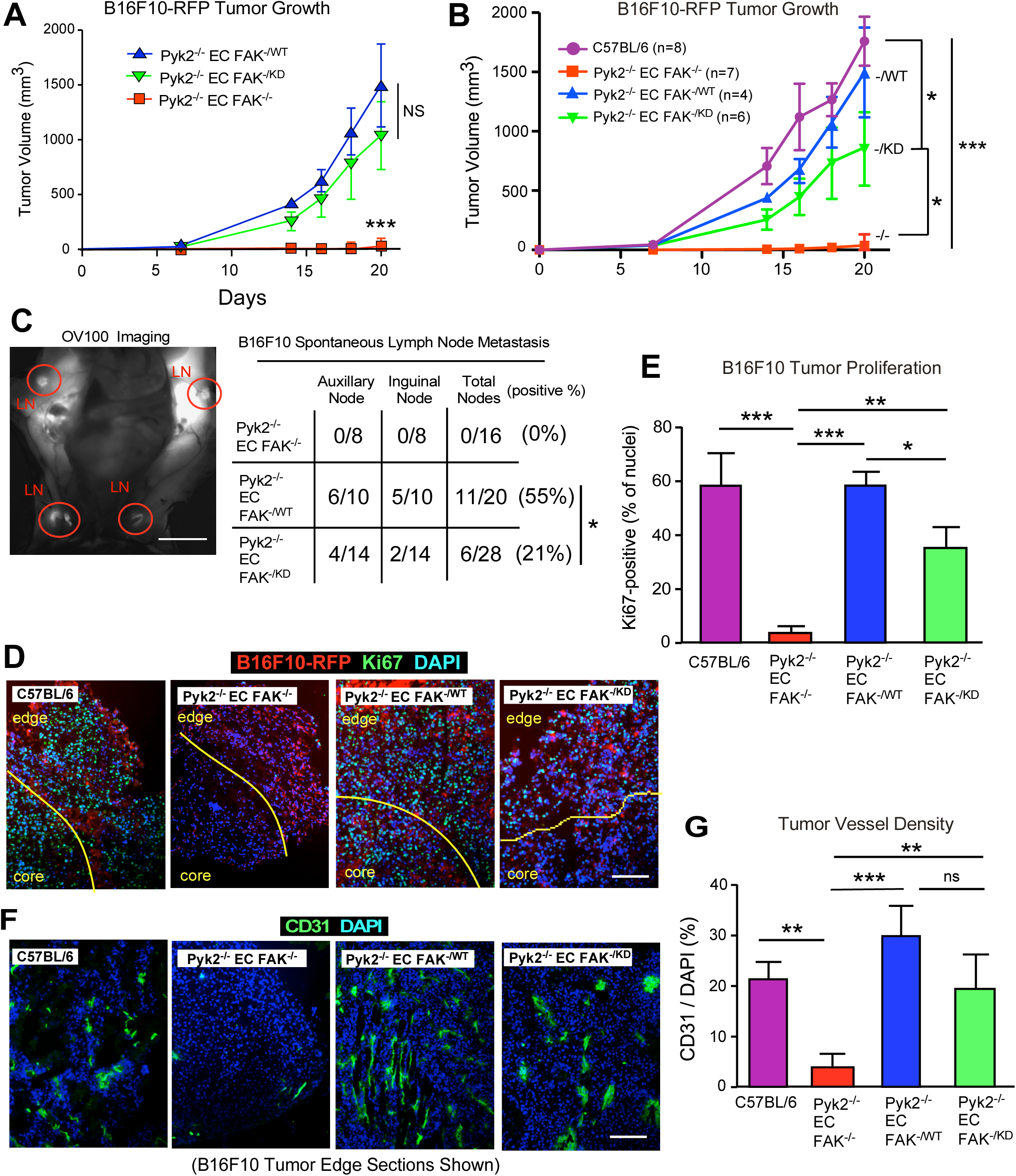
Hemizygous FAK KD expression in PYK2^-/-^ ECs supports tumor growth but limits metastasis. (**A**) Red fluorescent protein (RFP) labeled B16F10 melanoma subcutaneous tumor growth in tamoxifen-treated PYK2^-/-^ EC FAK^-/WT^ (blue triangles), PYK2^-/-^ EC FAK^-/KD^ (green triangles), and PYK2^-/-^ EC FAK^-/-^ (red squares) mice. Values are tumor volume means by caliper measurements (n = 8 mice per group, +/- SD). (**B**) B16F10-RFP tumor growth over 20 days in C57Bl6 (purple circles), PYK2^-/-^ EC FAK^-/WT^ (blue triangles), PYK2^-/-^ EC FAK^-/KD^ (green triangles), and PYK2^-/-^ EC FAK^-/-^ (red squares) mice. Values are tumor volume means +/- SD. (**C**) Spontaneous B16F10-RFP melanoma metastasis to auxiliary and inguinal lymph nodes at Day 21 as detected by *in situ* fluorescent detection. Values are from individual mice of indicated genotype (* *P* <0.05, T-test). (**D**) Representative sections of B16F10 tumors grown in tamoxifen-treated C57Bl6, PYK2^-/-^ EC FAK^-/-^, PYK2^-/-^ EC FAK^-/WT^, and PYK2^-/-^ EC FAK^-/KD^ mice. Frozen sections were fixed and stained with antibodies to Ki67 (green), DAPI (blue) and imaged by confocal microscopy (B16F10-RFP tumor cells, red). Scale is 50 µm. (**E**) Quantitation of B16F10-RFP proliferative index by Ki67 staining. Mean percentage of Ki67- and DAPl-positive cells (+/-SD, 10 sections from 3 tumors for each group. (**F**) Representative CD31 (green) and DAPI (blue) immunostaining of B16F10-RFP tumor sections from tamoxifen-treated normal C57BL6, PYK2^-/-^ EC FAK^-/-^, PYK2^-/-^ EC FAK^fl/WT^, or PYK2^-/-^ EC FAK^fl/KD^ mice. Scale is 20 μm. (G) Quantitation of CD31 vessel EC staining. Mean percentage of CD31 and DAPl-positive cells per field (+/-SD, 10 sections from 3 tumors for each group). Statistics for panels A, B, E, and G (ns, not significant, \**P* <0.05, ** *P* <0.01, *** *P* <0.001 ANOVA with Tukey’s post-hoc test).

Among PYK2^-/-^ EC FAK^-/-^ mice, the lack of primary B16F10 tumor formation can, unsurprisingly, impact tumor metastasis. However, overall B16F10 metastasis was also unexpectedly less among the PYK2^-/-^ EC FAK^-/KD^ mice compared to PYK2^-/-^ EC FAK^-/WT^ mice, at 21% and 55%, respectively (Figure 5C). Importantly, inhibition of B16F10 metastasis in PYK2^-/-^ EC FAK^-/KD^ mice occurred in experiments that showed no significant differences in tumor growth (Figure 5A). FAK-KD expression can limit VEGF-stimulated tumor metastasis via a reduced capacity to support tumor cell extravasation^22^. Since PYK2^-/-^ EC FAK^-/KD^ mice do not exhibit spontaneous vascular leakage as do PYK2^-/-^ EC FAK^-/-^ mice (Figure 4G), these results support the importance of intrinsic FAK activity in regulating EC permeability impacting tumor metastasis.

The lack of B16F10 tumor growth in PYK2^-/-^ EC FAK^-/-^ mice was also reflected in low levels of Ki67 staining associated with implanted tumors (Figures 5D and E). B16F10 tumors grown in C57Bl6 or PYK2^-/-^ EC FAK^-/WT^ mice exhibited high levels of Ki67 staining throughout the tumor edge and core regions (Figure 5D). In contrast, Ki67 staining was present in the edge region of B16F10 tumors grown in PYK2^-/-^ EC FAK^-/KD^ mice but generally absent within tumor cores (Figure 5D). This resulted in a significant decrease in overall Ki67 staining in tumor sections from PYK2^-/-^ EC FAK^-/KD^ compared to PYK2^-/-^ EC FAK^-/WT^ mice (Figure 5E). Upon further analysis of CD31 staining of B16F10 tumor edge regions, significantly fewer EC vessels were detected in tumors from PYK2^-/-^ EC FAK^-/-^ mice compared to other genotypes (Figure 5F and G). However, equivalent CD31 staining was detected in B16F10 tumor edge regions of C57Bl6, PYK2^-/-^ EC FAK^-/WT^, and PYK2^-/-^ EC FAK^-/KD^ mice. We speculate that the presence of CD31 ECs at the periphery but not the core of subcutaneous B16F10 tumors in PYK2^-/-^ EC FAK^-/KD^ mice may be related to observed vessel disorganization and EC sprouting alterations upon *in situ* labeling of tumor-recruited blood vessels determined by confocal microscopy in PYK2^-/-^ EC FAK^-/WT^ and PYK2^-/-^ EC FAK^-/KD^ mice (Figure 5 – supplemental figure 2). Taken together, these results support the notion that tumor angiogenesis is altered in PYK2^-/-^ EC FAK^-/KD^ mice, but loss of EC FAK activity remained sufficient to support tumor growth.

### Essential role for EC PYK2 and FAK expression in support of experimental tumor growth

To determine if the lack of B16F10 melanoma tumor growth in PYK2^-/-^ EC FAK^-/-^ mice extends to other syngeneic models, polyoma T antigen transformed Py8119 breast carcinoma cell growth as tumors was evaluated upon orthotopic mammary fat pad implantation (Figure 6 – figure supplement 1). By experimental Day 28, tumor volume was equivalent in PYK2^-/-^ EC FAK^-^

**Figure 6.**
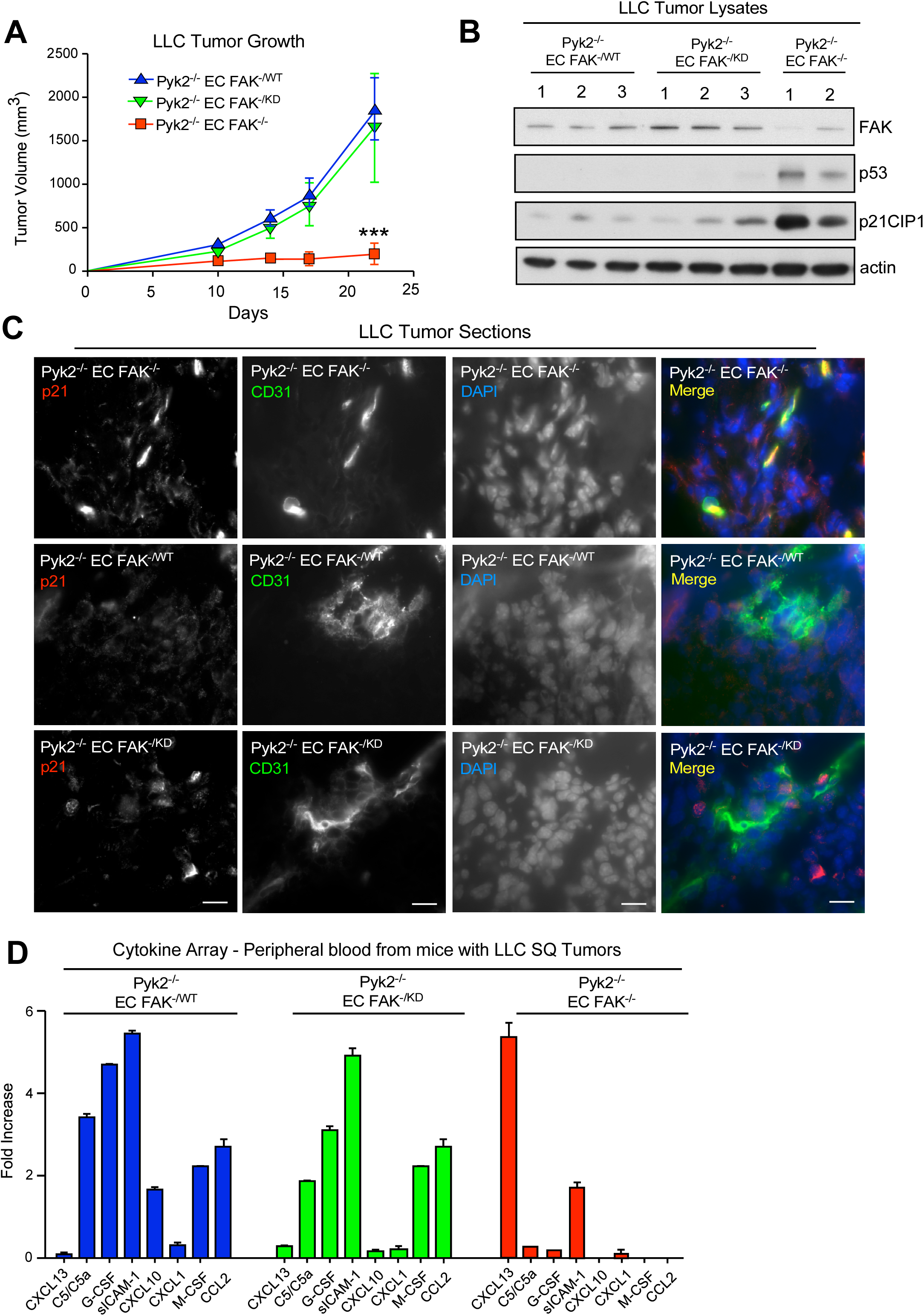
Inducible FAK loss but not prevention of FAK activity in ECs of PYK2^-/-^ mice boosts p53 activation to inhibit tumor growth. (**A**) Lewis lung carcinoma (LLC) subcutaneous tumor growth in tamoxifen-treated PYK2^-/-^ EC FAK^fl/WT^ (blue triangles), PYK2^-/-^ EC FAK^fl/KD^ (green triangles), and PYK2^-/-^ EC FAK^-/-^ (red squares) mice. Values are means of tumor volume +/- SD (n=8 mice per group, *** *P* <0.001 by ANOVA with Tukey’s post-hoc test) representing one of two independent experiments. (**B**) LLC tumor protein lysates were analyzed by immunoblotting for FAK, p53, p21CIP1, and actin in tamoxifen-treated mice of the indicated genotypes. Lanes represent lysates from individual tumors. (**C**) Representative p21CIP1 (red), CD31 (green), nuclei (DAPI, blue), and merged images of LLC frozen tissue sections from tamoxifen-treated PYK2^-/-^ EC FAK^-/-^, PYK2^-/-^ EC FAK^fl/WT^, and PYK2^-/-^ EC FAK^fl/KD^ mice. Yellow color denotes overlap staining. Scale is 20 μm. (**D**) Serum collected by venous puncture at euthanasia of LLC subcutaneous tumor-bearing mice of the indicated genotypes. Serum was pooled (n=4 mice per group), analyzed by the murine cytokine profiler array (R&D Systems), and values represent the mean of technical duplicates. Values are plotted as fold-increase compared to non-tumor bearing PYK2^-/-^ FAK^fl/fl^ Cre-mice.

^/WT^ and PYK2^-/-^ EC FAK^-/KD^ mice whereas only very small tumors formed in PYK2^-/-^ EC FAK^-/-^ mice. In a third syngeneic tumor model, Lewis lung carcinoma (LLC) cells were subcutaneously injected into mice of different FAK genotypes (Figure 6). LLC tumors grew equivalently in PYK2^-/-^ EC FAK^-/WT^ and PYK2^-/-^ EC FAK^-/KD^ mice but did not readily grow in PYK2^-/-^ EC FAK^-/-^ mice (Figure 6A). Analysis of tumor tissue lysates by immunoblotting revealed equivalent total FAK expression compared to actin in LLC tumors from PYK2^-/-^ EC FAK^-/WT^ and PYK2^-/-^ EC FAK^-/KD^ mice (Figure 6B). Surprisingly, only low levels of FAK were detected in residual LLC tumors from PYK2^-/-^ EC FAK^-/-^ mice. These lysates showed elevated levels of p53 and p21CIP1 proteins compared to actin (Figure 6B).

LLC tumor cells have a hyper-mutated genome, mutations in p53, and may be an unlikely source of elevated p53 and p21CIP1 proteins detected in lysates. To identify cells expressing p21CIP1, LLC tumors were evaluated by p21CIP1 and CD31 co-immunofluorescent staining (Figure 6C). p21CIP1 strongly co-stained with the few CD31 ECs present within LLC tumors from PYK2^-/-^ EC FAK^-/-^ mice. However, p21CIP1 staining did not localize with CD31 in LLC tumors grown in PYK2^-/-^ EC FAK^-/WT^ and PYK2^-/-^ EC FAK^-/KD^ mice where only low levels of background p21CIP1 staining was observed (Figure 6C).

Previous studies have shown that EC FAK inhibition can alter the production of cytokines^18^ and an ELISA-based array screen was performed on peripheral blood obtained from mice at the time of LLC tumor harvest (Figure 6D). A similar pattern of cytokines was detected in the blood of PYK2^-/-^ EC FAK^-/WT^ and PYK2^-/-^ EC FAK^-/KD^ mice, whereas a selective increase in CXCL13 and the decrease in the levels of other cytokines was detected in peripheral blood of tumor-bearing PYK2^-/-^ EC FAK^-/-^ mice (Figure 6D). These results support the notion that PYK2 and FAK loss within blood vessel ECs triggers both cell-intrinsic and cell-extrinsic changes resulting in a multi-factorial repressive or antagonistic environment for tumor growth.

### Inducible loss of FAK expression in primary PYK2^-/-^ mouse lung EC (MLEC) prevents survival in culture

As only a fraction of PYK2^-/-^ EC FAK^fl/fl^ Cre^+^ mice die after tamoxifen Cre activation *in vivo* (Table 1), we isolated primary MLECs by antibody-mediated affinity purification and verified rapid acetylated low-density lipoprotein uptake by these cells (Figure 7 – figure supplement 1). However, no viable PYK2^-/-^ FAK^-/-^ MLECs were obtained after tamoxifen pre-treatment of mice. As cell-permeable 4 hydroxy-tamoxifen (4-OHT) can be used to activate Cre-ERT in cells *ex vivo*, MLECs were isolated from various genotypes and immunoblotting performed 24 to 48 h after 4-OHT addition (Figures 7A-C). Accordingly, 4-OHT treatment reduced FAK levels in Cre^+^ but not Cre^-^ PYK2^-/-^ FAK^fl/fl^ MLECs and this was associated with increased p21CIP1 levels within 24 h (Figure 7B). Importantly, 48 h after 4-OHT treatment of PYK2^-/-^ FAK^-/WT^ and PYK2^-/-^ FAK^-/KD^ MLECs, no changes in p21CIP1 protein levels were detected (Figure 7C). Additionally, dual treatment of PYK2^-/-^ FAK^fl/WT^ MLECs with 4-OHT and small molecule ATP-dependent FAK-PYK2 inhibitor (VS4718) for up to 72 h, reduced MLEC FAK Y397 phosphorylation but did not result in increased p21CIP1 levels (Figure 7D). Together, these results show that combined loss of PYK2 and FAK expression but not inhibition of FAK activity within MLECs results in cell-intrinsic elevation of p21CIP1.

**Figure 7.**
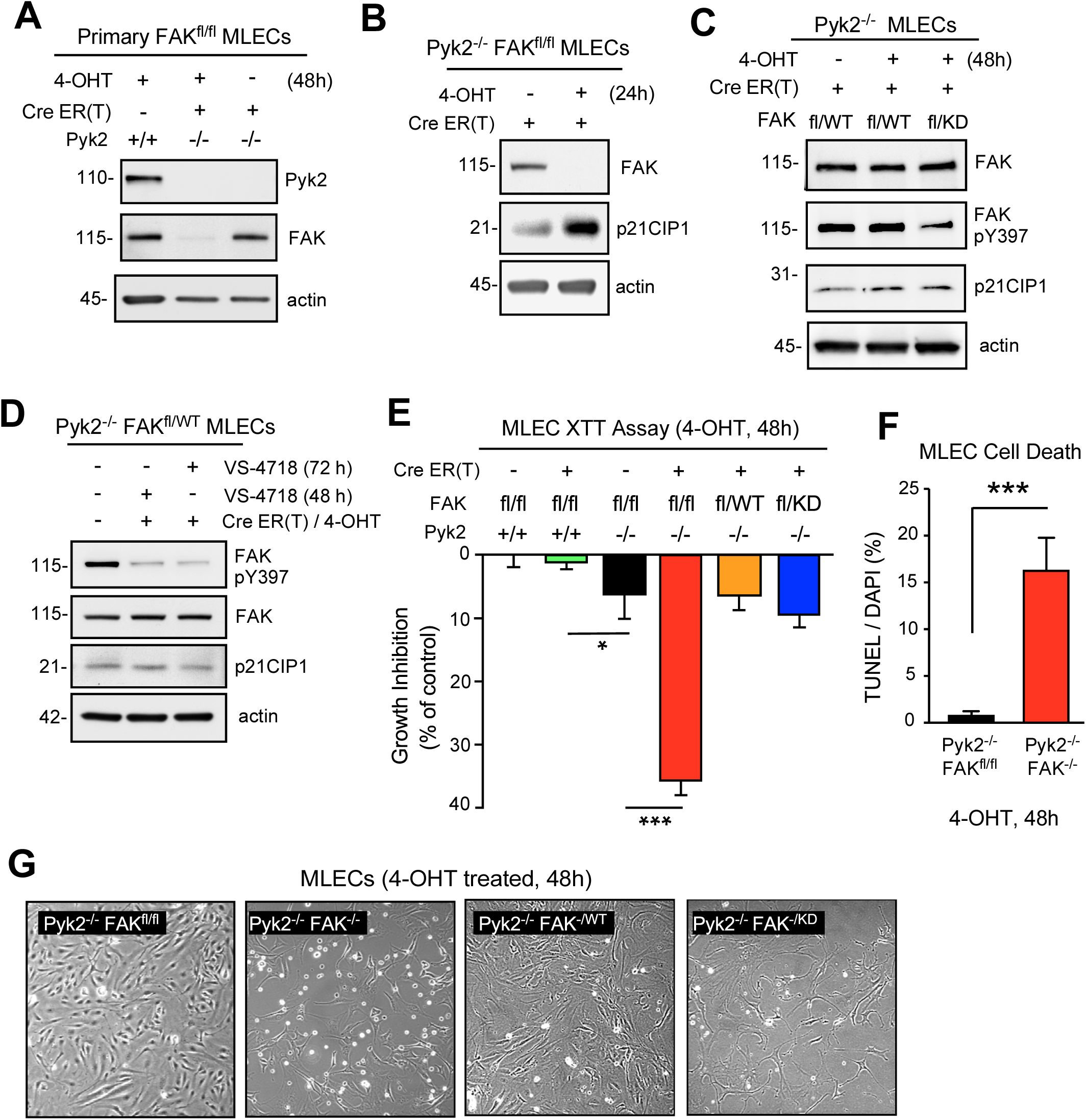
Inducible FAK loss but not prevention of intrinsic FAK activity limits PYK2^-/-^ MLEC survival in culture. (**A**) Primary MLECs of the indicated genotype before or after 4-hydroxy tamoxifen (4-OHT) addition (48 h) were immunoblotted for PYK2, FAK, and actin levels. (**B**) Cre+ PYK2^-/-^ FAK^fl/fl^ MLECs before or after 4-OHT addition (24 h) were immunoblotted for FAK, p21CIP1, and actin. (**C**) Cre+ MLECs of the indicated genotype before or after 4-OHT addition (48 h) were immunoblotted for FAK, pY397 FAK, p21CIP1, and actin. (**D**) PYK2^-/-^ FAK^fl/WT^ MLECs (+/- Cre) with or without pharmacological FAK inhibition (VS-4718, 1 µM) addition for 48 or 72 h were immunoblotted for FAK pY397, total FAK, p21CIP1, and actin. (**E**) Cell viability and proliferation (XTT tetrazolium salt assay) of primary MLECs of the indicated genotype after 4-OHT addition (48 h). Values are means +/- SD (n=4, two independent experiments, * *P* <0.05, *** *P* <0.001 by ANOVA) and are expressed as a percentage vehicle control-treated ECs of same genotype. (**F**) Quantification of 4-OHT stimulated cell death. Mean percentage of TUNEL- and DAPI-double positive MLECs per field. (+/- SD, *** *P* <0.001, T-test). (**G**) Phase contrast images of 4-OHT treated MLECs after 48 h. Magnification is 10X.

To determine if the induced loss of FAK on a PYK2^-/-^ background altered EC growth and metabolism *ex vivo*, equal numbers of Cre- and Cre+ MLECs of the indicated genotype were plated, 4-OHT was added, and cell metabolism measured by XTT assay (Figure 7E). Whereas Cre-mediated FAK inactivation on a PYK2^+/+^ cell background did not prevent MLEC growth, Cre+ PYK2^-/-^ FAK^-/-^ MLECs treated with 4-OHT exhibited significant growth inhibition (Figure 7E), increased cell death by TUNEL staining (Figure 7F), and cell rounding with loss of adhesion (Figure 7G). Notably, no significant differences were observed between 4-OHT-induced PYK2^-/-^ FAK^-/WT^ and PYK2^-/-^ FAK^-/KD^ MLEC growth (Figure 4E). Together, these results show that MLEC survival is compromised upon combined PYK2 and FAK loss and that this was associated with increased p21CIP1 expression.

### Loss of PYK2 and FAK in ovarian tumor cells triggers p53-dependent increased p21CIP1 and poly ADP-ribose polymerase (PARP) levels

Tumor cells with wildtype p53 often retain DNA damage responses compared to tumors harboring mutated p53^32,33^. ID8 cells are a commonly used murine ovarian tumor model that express wildtype p53^46,47^. KMF cells (*<u>K</u>ras*, *<u>M</u>yc*, and <u>F</u>AK-amplified) are derived from ID8 and grow readily in cell suspension^48^, a condition that normally triggers p53 activity^49^. Increased p21CIP1 levels were detected in suspended compared to adherent ID8 cell lysates (Figure 8A). Surprisingly, some increased levels of p53 and p21CIP1 were also detected in suspended compared to adherent KMF cell lysates (Figure 8A). By exome sequencing, KMF cells lack *Trp53* coding mutations, though, parallel RNA sequencing identifies altered Trp53 mRNA transcripts that contained introns and a thymidine insertion in the 3’ untranslated region which could potentially affect p53 protein translation (Figure 8B)^48^. Correspondingly, basal p53 protein levels in KMF are suppressed compared to ID8 cells (Figure 8A).

**Figure 8.**
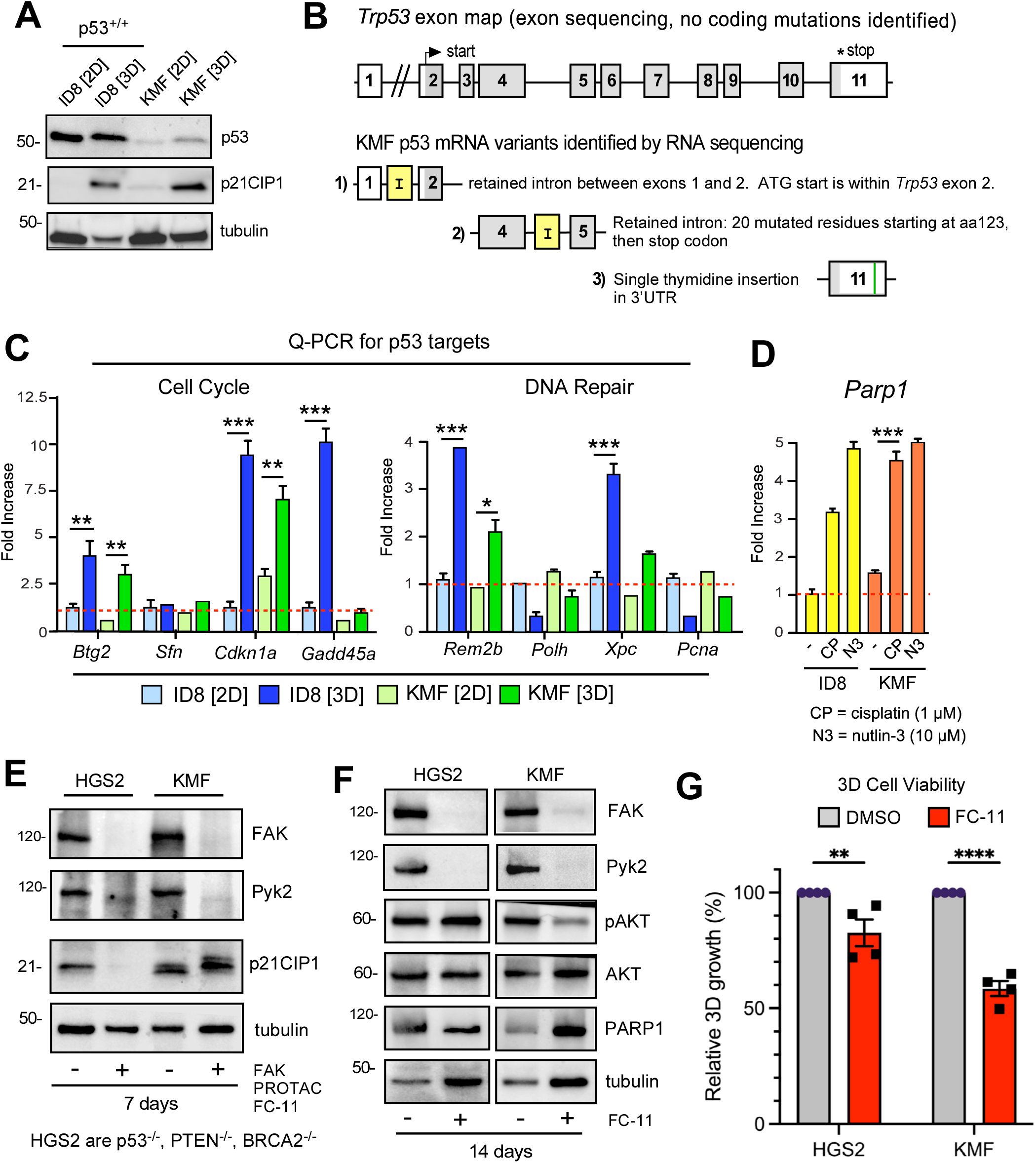
Loss of PYK2 and FAK in ovarian tumor cells triggers p53-dependent increased p21CIP1 and poly ADP-ribose polymerase (PARP) levels. (**A**) Equal numbers of ID8 and KMF ovarian tumor cells in were grown in adherent [2D] or suspended culture [3D] for 72 h and protein lysates were immunoblotted for p53, p21CIP1, and tubulin. (**B**) Schematic of *Trp53* exon map and identified p53 mRNA splicing alterations and single nucleotide polymorphism in the Trp53 exon 11 obtained from exon and RNA sequencing of KMF cells (NCBI Sequence Read Archive under accession number SRP194638 and Gene Expression Omnibus under the accession number GSE129099). (**C**) qRT-PCR of cell cycle or DNA repair mRNA targets regulated p53 activity using RNA collected from ID8 and KMF cells grown in 2D and 3D conditions for 72 h. (**D**) qRT-PCR analysis of *Parp1* mRNA levels in ID8 and KMF cells treated with cisplatin (1 µM) or nutlin-3 (10 µM) for 48 h. (**C** and **D**) Values are means +/- SEM (n=3, two independent experiments, * *P* <0.05, ** *P* <0.01, *** *P* <0.001 by ANOVA with Tukey post hoc test). (**E** and **F**) Lysates of HGS2 (p53^-/-^ PTEN^-/-^, BRCA2^-/-^) and KMF (partial wildtype p53 function) cells cultured in the presence of DMSO (control) or FAK PROTAC FC-11 (1 µM) for 7 or 14 days and immunoblotted for FAK, PYK2, p21CIP1, active Akt (pAKT), Akt, PARP, and tubulin. (**G**) Anchorage-independent HGS2 or KMF cell viability as measured by Alamar Blue in the presence of FC-11 (1 µM) or DMSO (control) after 72 h. Values are means +/- SEM of 3 technical replicates from 4 independent experiments (** P<0.01, **** P<0.0001 by 2-way ANOVA).

To measure potential p53-dependent transcriptional responses, ID8 and KMF cells were grown in 2D or anchorage-independent 3D conditions for 72 h, RNA isolated, and quantitative real-time PCR (qRT-PCR) analyses performed (Figure 8C). Increased cell cycle regulators *Btg2*, *Cdkn1a*, and *Gadd45a* as well as DNA repair targets *Rrm2b* and *Xpc* mRNA levels were detected in suspended versus adherent p53^+/+^ ID8 lysates. Additionally, *Btg2*, *Cdkn1a*, and *Rrm2b* mRNA levels were significantly increased in suspended KMF cell lysates (Figure 8C).

These qRT-PCR results are consistent with increased p21CIP1 levels (*Cdkn1a* mRNA) detected in lysates of 3D grown ID8 and KMF cells (Figure 8A). As chemotherapy stress can promote p53 activation, qRT-PCR was performed after cisplatin (1 µM, 24 h) treatment of ID8 and KMF cells to measure changes in mRNA levels of the DNA repair protein PARP1 in comparison to nutilin-3 treatment (10 µM), a small molecule activator of p53 (Figure 8D). Compared to DMSO-treated control cells, cisplatin elevated PARP1 mRNA in both ID8 and KMF cells. Notably, cisplatin and nutlin-3 induced equivalent levels of PARP1 mRNA in KMF cells, supporting the notion that KMF cells may retain partial p53 wildtype function. Compared to ID8 cells, KMF cells express high levels of FAK and Pyk2, readily proliferate in suspension conditions, and exhibit no growth defects in 2D culture upon FAK knockout^48^.

PROTACs (proteolysis targeting chimeras) have been developed targeting FAK and PYK2 and these molecules work by inducing selective intracellular proteolysis^8^. The murine HGS2 ovarian cancer model, derived from p53^-/-^, PTEN^-/-^, and BRCA2^-/-^ transgenic mice, expresses high levels of FAK and PYK2 like KMF cells (Figure 8E). To determine if PROTAC knockdown of both PYK2 and FAK could trigger differential p53-associated changes in KMF (partial p53 function) compared to HGS2 (p53^-/-^), cells were treated with DMSO (control) or FAK PROTAC FC-11 (1 µM) and lysates were analyzed by immunoblotting after 7 or 14 days (Figure 8E and F). FAK PROTAC FC-11 is derived from the FAK inhibitor PF-562271 which inhibits both FAK and PYK2 with nanomolar affinity^50^.

After 7 days, FAK and PYK2 proteins were no longer detected in FC-11-treated HGS2 and KMF cell lysates (Figure 8E). Interestingly, whereas p21CIP1 protein was present in DMSO control cells, it was not detected in FC-11-treated HGS2 cell lysates after 7 days. However, in KMF cells, p21CIP1 was detected in control and FC-11-treated cell lysates where p21CIP1 exhibited a slightly increased size by gel electrophoresis (Figure 8E). As HGS2 and KMF cell number increased after 7 days in the presence of FC-11 (data not shown), the cells were split, equal numbers were plated in the presence of FC-11 or DMSO for another 7 days, and lysates analyzed by immunoblotting (Figure 8F). After 14 days, FAK and PYK2 levels remained undetectable with FC-11 treatment, and HGS2 cells exhibited elevated levels of activated Akt compared to KMF, possibly related to PTEN inactivation in HGS2 cells. In comparison, KMF cells showed higher PARP1 levels than HGS2 after 14 days of FC-11 treatment (Figure 8F). Elevated PARP1 may represent one element of a p53-dependent DNA damage response, with DNA repair occurring over time in FC-11-treated KMF cells compared to p53^-/-^ HGS2 cells. To determine if PROTAC FC-11 degradation of FAK and Pyk2 may impact anchorage independent growth, DMSO- or PROTAC FC-11-treated HGS2 and KMF cells were analyzed after 72 h by an Alamar Blue metabolic assay (Figure 8G). Notably, FC-11 treatment resulted in a greater inhibition of KMF cell viability compared to HGS2 cell response. Altogether, our results using FAK PROTAC FC-11 support the notion that loss of FAK-PYK2 expression triggers a cell intrinsic stress response that can also boost p53-associated signals in tumor cells.

## Discussion

FAK possesses a similar domain structure and regulatory phosphorylation sites as PYK2, and thus FAK-PYK2 have been hypothesized to act in parallel or, in some cases, as redundant signaling proteins^8^. PYK2 knockout mice are viable and fertile^26^, while EC-specific FAK knockout or genetic inactivation of EC FAK activity results in early developmental lethality^14,25^, clearly illustrating that some functions of FAK cannot be efficiently duplicated by PYK2. Indeed, EC FAK loss in adult mice can result in variable phenotypes^17,38,39^. The limited phenotypes of PYK2^-/-^ mice support the notion that FAK may play important compensatory roles in PYK2^-/-^ cells. Herein, we created and characterized the phenotypes associated with inducible FAK loss of expression or inducible genetic suppression of FAK activity within adult mouse ECs on PYK2^+/+^ or PYK2^-/-^ mouse backgrounds. PYK2^-/-^ EC FAK^-/-^ mice exhibit elevated vascular leakage and a strong anti-tumor phenotype using three different syngeneic models. This PYK2^-/-^ EC FAK^-/-^ anti-tumor phenotype was associated with abortive EC vessel sprouting, EC-increased p53 and p21CIP1 protein levels, and alterations in peripheral blood cytokines. The induced loss of EC FAK expression in PYK2^-/-^ mice was associated with p53 activation *in vivo* and *in vitro* (Figure 9).

**Figure 9.**
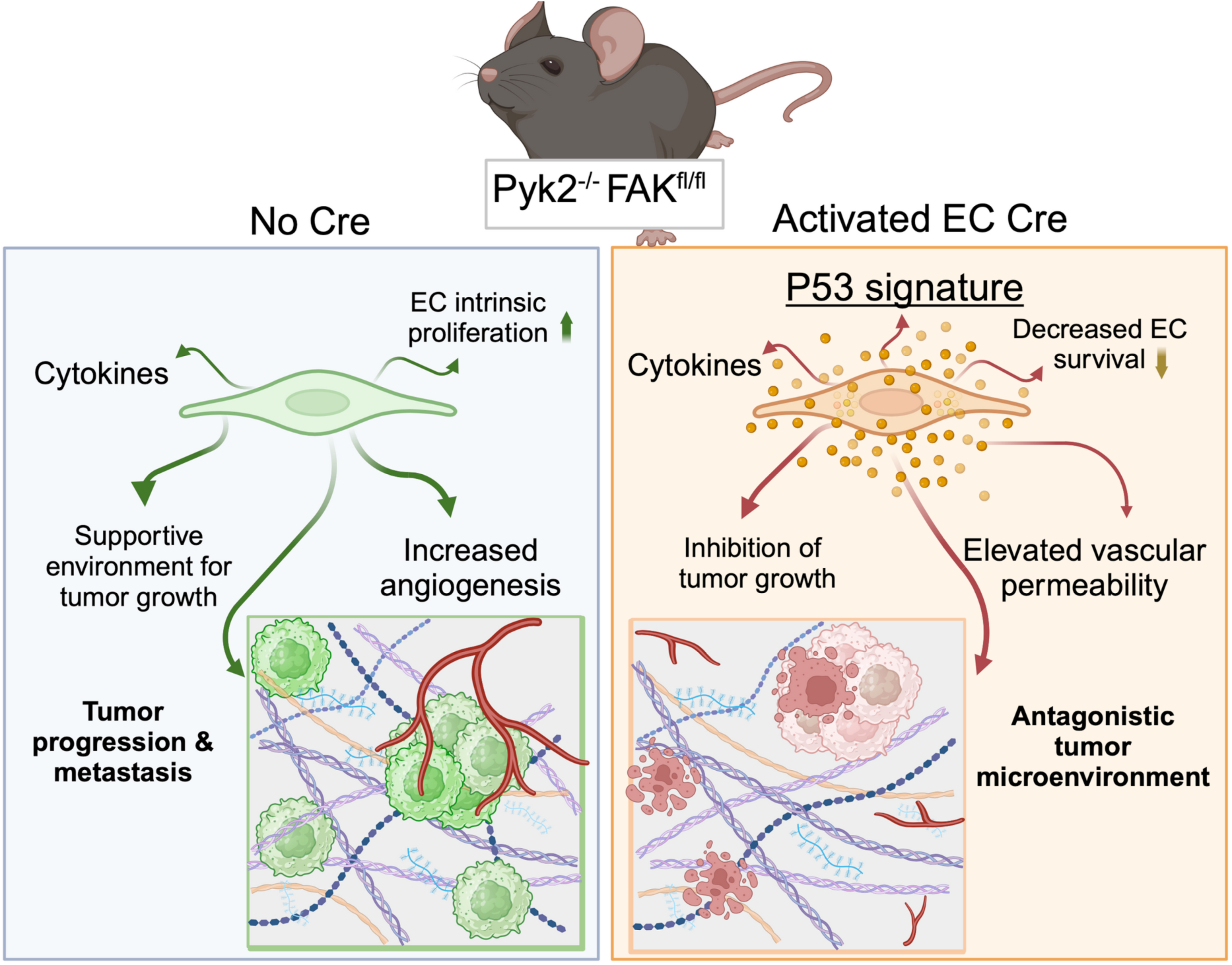
Schematic of inducible EC FAK knockout in Pyk2^-/-^ mice with negative p53-associated effects on tumor growth. Tamoxifen-activated and EC-enriched Cre-recombinase expression was used to inactivate FAK expression creating PYK2^-/-^ EC FAK^-/-^ mice and comparisons were made to Cre-negative PYK2^-/-^ EC FAK^fl/fl^ littermates that exhibited normal EC function and a supportive environment for tumor growth and metastasis like PYK2^+/+^ mice. In contrast, a hostile tumor microenvironment is created in PYK2^-/-^ EC FAK^-/-^ mice with p53 tumor suppressor activation, increased p21CIP1 cell cycle inhibitory protein expression, and decreased primary EC survival in culture. Loss of PYK2 and FAK in ECs results in loss of vascular integrity and the inhibition of tumor growth. Combined with results using of PROTAC degraders of FAK and PYK2 to trigger p53-associated signaling in ovarian tumor cells, loss of FAK-PYK2 expression triggers a cell intrinsic stress response that may also boost p53-associated cell death signals in tumor cells. Created in BioRender. Schlaepfer, D. (2024) BioRender.com/r33u123

Growing tumors are surrounded by a mixed microenvironment of fibroblasts, endothelial-lined blood vessels, immune cells, and changes in the composition of growth factors and extracellular matrix^51^. Crosstalk between normal stromal and tumor cells controls tumor progression and spread. Although p53 tumor suppressor expression can function as a cell-intrinsic barrier to tumor progression in part by promoting the expression of cell cycle inhibitory proteins such as p21CIP1, and DNA repair proteins such as PARP1^32,33^, studies have also uncovered cell extrinsic roles for p53 in creating an anti-tumor microenvironment^34^. These p53 effects can be mediated through alterations in matrix protein secretion or gene expression driving elevated cytokine factors altering immune surveillance and cell senescence. As previous studies have shown that EC FAK signaling supported tumor survival from chemotherapy by EC cytokine production^18^, our findings of altered blood cytokine levels in PYK2^-/-^ EC FAK^-/-^ mice bearing tumors support the notion that EC vessels lacking FAK-PYK2 expression may act to create an antagonistic or anti-tumorigenic microenvironment.

The ability of FAK to suppress p53 was first observed during development^25^. FAK^-/-^ mouse embryo fibroblasts (MEFs) were resistant to standard embryo outgrowth methods, and this was bypassed upon FAK inactivation on either a p53^-/-^ or p21^-/-^ mouse background^30^. In contrast, PYK2^-/-^ MEFs exhibit no such p53- or p21-dependent growth restrictions. Although Pyk2 knockdown in FAK^-/-^ p21^-/-^ MEFs resulted in elevated p53 activation and growth arrest^31^, we speculate that p53 regulation may be more tightly linked to changes in FAK expression as reflected by our results using tamoxifen-inducible PYK2^-/-^ EC FAK^-/-^ mice. While more focused studies are required, we speculate that vessel-associated p53 activation in PYK2^-/-^ EC FAK^-/-^ mice may represent extrinsic cell stress associated with vascular barrier failure, tissue edema, and elevated fibrosis. However, as primary PYK2^-/-^ FAK^-/-^ MLECs exhibit induced p21CIP1 levels and cell death with 24-48 h upon induced FAK loss in culture, there is also intrinsic cell stress generated by FAK-PYK2 loss that supports rapid p53 activation. This “keeper of the guardian” (ie., p53) role for FAK is consistent with findings showing that FAK exhibits nuclear accumulation in various cell types upon treatment with drugs that prevent nuclear protein export^30,52^.

Genomic and pharmacologic approaches can sometimes differ in outcome. Herein, PROTAC-mediated knockdown of FAK and PYK2 also resulted in p53-associated changes in cell cycle inhibitors such as p21CIP1 and DNA repair proteins such as PARP1. Our results underscore the potential importance of p53 regulation by loss of FAK-PYK2 expression impacting tumor growth, as was implicated in studies targeting FAK knockdown in p53^+/+^ mesothelioma^53^. Currently, small molecule FAK ATP-competitive inhibitors are being tested clinically in several combinatorial Phase II and Phase III trials^8^. The generation of PYK2^-/-^ EC FAK^-/WT^ and PYK2^-/-^ EC FAK^-/KD^ mice with tamoxifen-induced hemizygous wildtype or kinase-deficient FAK expression revealed that the scaffolding role for EC FAK and not kinase activity was sufficient to support primary tumor growth, but not metastasis. As pharmacological FAK inhibition can prevent tumor cell motility, vascular permeability growth factor and matrix metalloproteinase expression^54,55^, our results with PYK2^-/-^ EC FAK^-/KD^ mice suggest that inhibition of FAK-PYK2 activity may be a potent mechanism acting on tumors and the vasculature to control cancer spread.

Collectively, our studies begin to delineate the specific effects of FAK inhibition within ECs and tumor cells. Genetically, loss of Pyk2-FAK expression is not equivalent to the inhibition of Pyk2-FAK activity. Moreover, there may be biological differences in drugs that selectively target FAK and those that co-target FAK and Pyk2. Lastly, as PROTAC degraders advance in clinical testing^56^, it will be interesting to determine whether combined FAK-PYK2 targeting may be an approach to actively induce a p53-associated anti-tumor response.

## Materials and Methods

### Mice

Mice with loxP sites flanking FAK exon 3 (FAK^fl/fl^) and a tamoxifen-inducible Cre-estrogen receptor tamoxifen (ERT) fusion protein (Cre ERT) transgene downstream of the 5′ endothelial enhancer of the stem cell leukemia locus^39^ were crossed with PYK2 knockout (PYK2^-/-^) mice^26^ to generate PYK2^-/-^ FAK^fl/fl^ Cre-ERT mice. PYK2^-/-^ mice were crossed with heterozygous mice containing a kinase-deficient (K454R) mutation (FAK^KD/WT^) within FAK exon 15^14^ to generate PYK2^-/-^ FAK^WT/KD^ mice. Founder mice were backcrossed to C57Bl6J for ten generations, transgenic mouse background purity (>97% B6J) was verified by single nucleotide polymorphism genome scanning analysis (Jax Services), and age-matched littermates were used for all experiments. Mouse progenies were identified by genotyping using PCR as described^14,26,39^. At 6 weeks of age, a combination of males and females PYK2^-/-^ FAK^fl/fl^ Cre^+^, PYK2^-/-^ FAK^fl/WT^ Cre^+^ and PYK2^-/-^ FAK^fl/KD^ Cre^+^ (and equal numbers of Cre-negative littermates as controls) mice were fed tamoxifen-chow (Envigo, TD.55125) for 3 weeks and rested for 1-2 weeks prior to experimental analyses. In some experiments, mice were treated with 2 mg tamoxifen (Sigma-Aldrich) every 2 d (IP injection in corn oil) for 2 weeks and then rested 2 weeks prior to analyses. All mouse procedures were reviewed [protocol S07731] and approved by The University of California San Diego Institutional Animal Care and Use Committee.

### Antibodies and reagents

Antibody to FAK (clone 4.47) was from Millipore Sigma. Antibody to PYK2 (clone 5E2), to GAPDH (clone 14C10), to PARP (clone 46D11), and γ-H2AX pSer139 polyclonal antibodies (#2577) were from Cell Signaling Technology. CD31 (PECAM-1) antibody (clone MEC 13.3) and to CD102 (ICAM-2) antibody (clone 3C4) were from BD Pharmingen. P53 polyclonal antibodies (FL-393) and antibody to p21CIP1 (clone F-5) were from Santa Cruz Biotechnology. Anti-FAK pY576 polyclonal antibodies (ab226847), antibody to FAK pY397 (clone EP2160Y), and antibody to α-SMA antibody (clone 1A4) were from Abcam. HP1γ antibody (clone 2MOD-1G6) and p16INK4 antibody (clone 1E12E10) were from Invitrogen. Actin antibody was from (clone AC-74) was from Sigma-Aldrich. VS-4718 FAK inhibitor and Nutlin-3 were from MedChemExpress and FAK PROTAC FC-11 was from Tocris Inc.

### Immunohistochemical staining

Paraffin-embedded heart and lung tissue were sectioned and processed for hematoxylin and eosin (H&E) staining and hematoxylin-Van Gieson staining (ATOM Scientific). Tissue sections were deparaffinized, rehydrated, processed for antigen retrieval, and quenched for peroxidase activity. Sections were blocked (PBS, 1% BSA, and 0.1% Triton X-100) for 45 min at RT and incubated with anti-p53 (1:50) and anti-p21Cip1 (1:50) overnight followed by biotinylated goat anti–rabbit IgG (1:300) for 30 min, and antibody binding was visualized with diaminobenzidine and counterstained with Methyl Green (Vectastain ABC Elite; Vector Laboratories). Images were acquired using upright (Olympus BX43) microscope with UPlanSApo 4X (NA 0.16), UPlanSApo 10X (NA 0.40), and UPlanSApo 20X (NA 0.75) objectives, and a digital color camera (Olympus SC100). Files were cropped and contrast adjusted using Photoshop (Adobe).

### *In vivo* vascular permeability

Mice were injected 100 μl Evans blue (Sigma, 20 mg/ml) via tail vein. After 30 min (heart) or 3 h (lungs), mice were sacrificed, perfused with 15 ml of saline, lungs and heart removed, photographed and tissue homogenized in 500µl formamide. Evans blue dye was extracted from tissues by incubation at 60 °C overnight. The tissue was pelleted by centrifugation (12,000 *x g* for 30 min), and the concentration of dye extracted in the supernatant was determined by spectroscopy (610 nm).

### Immunoblotting

Cells in culture or tissues collected from mice were washed with cold PBS, whole cell protein lysates were made by RIPA Lysis Buffer (Pierce) addition, and lysates were clarified by centrifugation (16,000 x g, 5 min). Complete™ Mini ETDA-free Protease inhibitor cocktail (Millipore Sigma) and PhoSTOP™ phosphatase inhibitor cocktail (Millipore Sigma) were added prior to use. Total protein levels were determined using a bicinchoninic acid assay (Pierce), 25 µg of protein were resolved on Mini-Protean TGX precast gels (4-15% Tris/Glycine gel, BioRad), and transferred to polyvinylidene difluoride membranes using a TransBlot Turbo (BioRad). Immunoreactive protein bands were detected using HRP-conjugated anti-mouse or anti-rabbit antibodies with Clarity Western ECL (enhanced chemiluminescence) substrate reagent and visualized using a ChemiDoc™ Touch Imaging System (BioRad).

### Tissue and cell staining

Frozen heart and lung sections in Optimal Cutting Temperature (OCT) compound were prepared (7 μm, Leica CM1950), fixed in cold acetone (10 min), rehydrated in PBS containing 0.5% BSA (5 min), and blocked with 1.25% normal goat serum in PBS (30 min at RT). Where indicated, samples were incubated with antibodies to FAK (1:50), FAK pY576 (1:100), CD31 (1:300), p21Cip1 (1:100), α-SMA (1:200), or γ-H2AX pS139 (1:200) overnight at 4°C followed by Alexa Fluor-488 and Alexa Fluor-594 secondary antibodies (Thermo Fisher 1:500, 30 min at RT), and DAPI (300 nM, D1306 Thermo Fisher) staining for 5 min. Imaging was performed sequentially using an Olympus IX81 spinning disk confocal microscope with zero drift compensation focus control, UPlanAPO 20x (NA 0.70) air, UPlanFL 40x (NA/1.30) oil, and PlanApo 60x (NA 1.42) oil objectives, and Hamamatsu OrcaER camera controlled by Slidebook software. Files were cropped, pseudo-colored, and contrast-adjusted using Adobe Photoshop. Degree of association exhibited by patterns of fluorescence was measured on a pixel-by-pixel basis and calculated as a Pearson’s correlation coefficient using the “measure correlations” module (Cell Profiler, v2.0, Broad Institute). A value of 0 indicates no overlap and a value of 1 corresponds to 100% colocalization.

For tissues, TUNEL (terminal deoxynucleotidyl transferase dUTP nick end labeling) staining, frozen heart and lung sections were fixed in 4% paraformaldehyde, permeabilized with 0.1% Triton X-100 (3 min), and incubated with anti-CD31 (1:300) overnight at 4°C followed by Alexa Fluor-594 secondary antibodies (Thermo Fisher 1:500, 30 min at RT) as recommended (*In Situ* Cell Death Detection Kit, Millipore Sigma). For cells, MLECs were grown to confluency on 0.1% gelatin coated glass coverslips, fixed in 4% paraformaldehyde (30 min at RT), permeabilized with 0.1% Triton X-100 (3 min) and incubated with antibodies as above. Ten TUNEL-stained images from 3 coverslips were acquired. Only TUNEL-positive colocalized with DAPI cells were enumerated using ImageJ.

For measurement of Ki67 proliferative marker staining, B16F10-RFP tumors were excised, frozen in OCT, sectioned, fixed in 4% paraformaldehyde, processed for antigen retrieval using citrate buffer (pH 6.0), and incubated with antibodies to Ki67 (1:100) overnight at 4°C followed by Alexa Fluor-488 secondary antibodies (Thermo Fisher 1:500, 30 min at RT), and DAPI (300 nM, Thermo Fisher) staining for 5 min. Sequential imaging using an Olympus IX81 spinning disk confocal microscope was performed as above.

### Tumor growth

Murine B16F10 melanoma (CRL-6475), Lewis lung carcinoma (LLC, CRL-1642), and Py8119 (CRL-3278) breast adenocarcinoma cells were purchased from ATCC. RFP lentivirus (Cell BioLabs) was used to make B16F10-RFP by puromycin selection and cell enrichment by flow cytometry sorting. C57Bl6 mice flanks were shaved and 1×10^6^ B16F10-RFP or LLC cells mixed with growth factor reduced Matrigel (200 µl) were injected subcutaneously, or Py8119 breast carcinoma cells were injected into the T4 mammary fat pad of 11 wk old tamoxifen pre-treated mice. Tumors were measured every 3-4 d with digital vernier calipers and tumor volume (mm^3^) was calculated using the formula: V = L x W^2^/2 (L = length, mm; W = width, mm). Body weight was measured weekly. Lungs, spleen, and primary tumors were surgically removed and weighed. Tumors sections were homogenized in protein lysis buffer for immunoblotting or placed in OCT (Tissue Tek), frozen in liquid nitrogen, thin sectioned (7 mm) using a cryostat (Leica 3050S), and mounted onto glass slides. To determine spontaneous B16F10-RFP metastasis, mice were euthanized, body cavity surgically opened and in situ fluorescence acquired using an Olympus OV100 small animal imaging system. A common threshold of RFP fluorescence was set for all OV100 acquisition and lymph node positivity for tumors was verified by imaging nodes after dissection.

### In vivo angiogenesis

The Matrigel assay was performed to assess *in vivo* angiogenesis. In brief, mice were injected subcutaneously on left and right flanks with 5×10^6^ B16F10-RFP tumor cells in 400 µl of growth factor-reduced Matrigel (Corning, 354230). After 7 d, mice were injected intravenously with 20 μg Fluorescein-labeled Griffonia (Bandeiraea) Simplicifolia Lectin I (FL-1101, Vector Labs) that labels vessel ECs. After 30 min, tumor-matrigel plugs were removed, photographed, fixed in paraformaldehyde, and images were acquired using laser scanning confocal microscopy (Nikon C1si) with Plan Apo 10X air (NA 0.45), Plan Apo 20X air (NA 0.75), and Plan Apo 60X (NA 1.40) oil objective lenses (Nikon) and post-processed for spectral illumination.

### Blood cytokines

Tumor-bearing mice were exsanguinated at time of euthanasia by trans cardiac puncture. Blood was allowed to clot at RT (30 min) and serum was collected after centrifugation (2000xg). 5 ml of serum (collected and combined from n=4 mice) was used to probe the Mouse Cytokine Proteome Profiler Array (R&D Systems) according to the manufacturer’s instructions.

### Endothelial cells

Primary MLECs were isolated from lung tissue by magnetic bead anti-CD31 and anti-CD102 binding as described^21^. Isolated ECs were verified for acetylated low density lipoprotein uptake and for CD31 plus ICAM-2 surface expression by flow cytometry as previously described^21^. MLECs were treated with adenoviral Cre (Cell Biolabs) or 4-hydroxy-tamoxifen (4-OHT, 2 µM for 24 to 48 h) to promote floxed FAK excision and used at passages <5 without immortalization. MLECs were maintained in EBM-2 basal EC medium (Lonza) with microvascular supplementation (BulletKit; Lonza) and 10% FBS on plates coated with 0.2% gelatin and 10 µg/ml rat tail collagen I (EMD Millipore).

### Ovarian tumor cells

KMF ovarian tumors cells were propagated as described^48^. HGS2 (p53^-/-^, PTEN-/-, and BRCA2^-/-^) ovarian tumor cells were obtained from Ximbio.com (#160538)^57^ and grown in Advanced DMEM/F-12 media (Thermo), 4% fetal bovine serum, 1X insulin-transferrin-selenium, 100 ng hydrocortisone, 20 ng/ml EGF, 1X penicillin-streptomycin and L-glutamine. For cell assays, 1×10^6^ cells were plated in 10 cm cell culture dishes in the presence of 1 µM FC-11 or DMSO and after 7 days, FC-11 treated HGS2 and KMF cells were replated under the same conditions for an additional 7 days prior to analysis.

### Cell viability and proliferation

Cells were seeded in a 96-well plate (1×10^5^ cells/well) in 100 μl of culture medium and incubated overnight. The following day, 4-OHT was added and after 48 h, cell metabolic activity was measured using an XTT Cell Proliferation Assay Kit (Cell Signaling Tec.). Absorbance values was measured at 450 nm by a plate reader (Tecan Spark) and represent the mean of triplicate points for all experiments. To measure 3D anchorage independent growth, KMF or HGS2 cells (5000 cells/well) were seeded on Ultra Low Attachment Surface 96 well plate (Corning, #3474) in 100µl of complete growth media. DMSO as control or FC-11 (1 µM) was added in 50 µl of complete media after 24 h. Alamar Blue Reagent (Invitrogen) at 10% of total volume (15 µl) was added 72 h following FC-11 treatment, cells were incubated at 37°C in 5% CO2 for 24 h, and absorbance at A570/600nm determined by plate reader (SpectraMax ID3, Molecular Devices).

### Quantitative RT-PCR

Total RNAs were extracted using PureLink™ RNA Mini Kit (Thermo) and cDNA prepared using the High-Capacity cDNA Reverse Transcription Kit (Thermo) from 1 µg total RNA. Target transcripts were amplified using a LightCycler 480 (Roche Applied Science), Premix Ex Taq probe qPCR Kit, iTaq™ Universal SYBR® Green Supermix (Bio-rad) with cDNA templates and primers according to manufacturer instructions. Target gene expression was normalized to 40S ribosomal protein S17 (RS17) as a housekeeping gene control. Transcript levels were calculated using the ΔΔCT (cycle threshold) method. Primers used were: Btg2 (F) CGCACTGACCGATCATTACAA. Btg2 (R) GGATCAACCCACAGGGTCAG, Cdkn1a (F) GCAGACCAGCCTGACAGATTT, Cdkn1a (R) TGGGCACTTCAGGGTTTTCT, Gadd45a (F) TGGTGACGAACCCACATTCA, Gadd45a (R) CGGGAGATTAATCACGGGCA, Parp1 (F) GGGCAAGCACAGTGTCAAAG, Parp1 (R) TGTCGTTGACACCAGATGGG, Pcna (F) AAAGATGCCGTCGGGTGAAT, Pcna (R) TCTATGGTTACCGCCTCCTCT, Polh (F) CCTCGCTATGACGCTCACAA, Polh (R) AGGAGGGGACCACTCAGTTT, Rrm2b (F) CTGTTCAAATCGAGCAGGAGT, Rrm2b (R) GCCATGACTGCAAATCGCTG, Sfn (F) ACAGGCCGAACGGTATGAAG, Sfn (R) GTACTCTTTCACCTCGGGGC, Xpc (F) ATTCCAGGGATTGCGTGCAT, and Xpc (R) TCCCAAACTCATTCCGAGGC

## Statistics

Differences between groups (ANOVA, one-way analysis of variance with Tukey’s post hoc) and differences between pairs of data (unpaired two-tailed Student’s *t* test) were analyzed using Prism (v9; GraphPad Software) and values of <0.05 were considered significant.

## Funding

This work was funded by National Institutes of Health grants to D. Schlaepfer and D. Stupack (R01CA247562 and R01CA254342), a V Foundation Translational Grant to D. Schlaepfer (T2023-018), and by CCSG support to the UCSD Moores Cancer Center (P30CA023100) for the flow cytometry and microscopy shared resources. M. Ojalill was supported in part by a Sigrid Juselius Foundation Award.

## Author Contributions

Xiao Lei Chen - conceptualization, data collection and analysis, formal analysis, and editing.

Marjaana Ojalill - data collection and analysis, visualization, formal analysis, and editing.

Christine Jean - conceptualization, data collection and analysis, formal analysis, and editing.

Isabelle Tancioni - data collection and analysis, visualization, and formal analysis.

Antonia Boyer - data collection and analysis.

Shulin Jiang - data collection and analysis.

Duygu Ozmadenci- data collection and analysis.

Sean Uryu - data collection and analysis.

David Tarin - visualization and methodology.

Joseph Schlessinger – materials.

Dwayne G. Stupack – conceptualization, supervision, visualization, formal analysis, and writing-review and editing.

David D. Schlaepfer – conceptualization, supervision, visualization, formal analysis, methodology, writing-original, project administration, writing-review and editing.

## Conflict of Interest

Dwayne Stupack is a consultant for Amplia Therapeutics Limited.

Other authors declare that they have no conflicts of interest.

## List of Abbreviations

4-OHT: 4-hydroxytamoxifen
CRE-ERT: Cre recombinase mutant estrogen receptor (ERT) ligand-binding domain fusion
DAPI: 4′,6-diamidino-2-phenylindole
EC: endothelial cell
FAK: focal adhesion kinase
fl: floxed
KD: kinase defective
KO: knockout
LLC: Lewis lung carcinoma
MLECs: murine lung endothelial cells
OCT: optimal cutting temperature
PARP1: Poly [ADP-ribose] polymerase 1
p21CIP1: cyclin-dependent kinase inhibitor 1
PROTAC: proteolysis targeting chimera
qRT-PCR: quantitative real-time reverse transcription polymerase chain reaction
RFP: red fluorescent protein
SCL: stem cell leukemia
SMA: smooth muscle actin
TUNEL: terminal deoxynucleotidyl transferase dUTP nick end labeling
VEGF: vascular endothelial growth factor
WT: wildtype

