## Supplementary files for "Inducible FAK Deletion but not FAK Inhibition in Endothelial Cells Activates p53 to Suppress Tumor Growth in PYK2-null Mice"

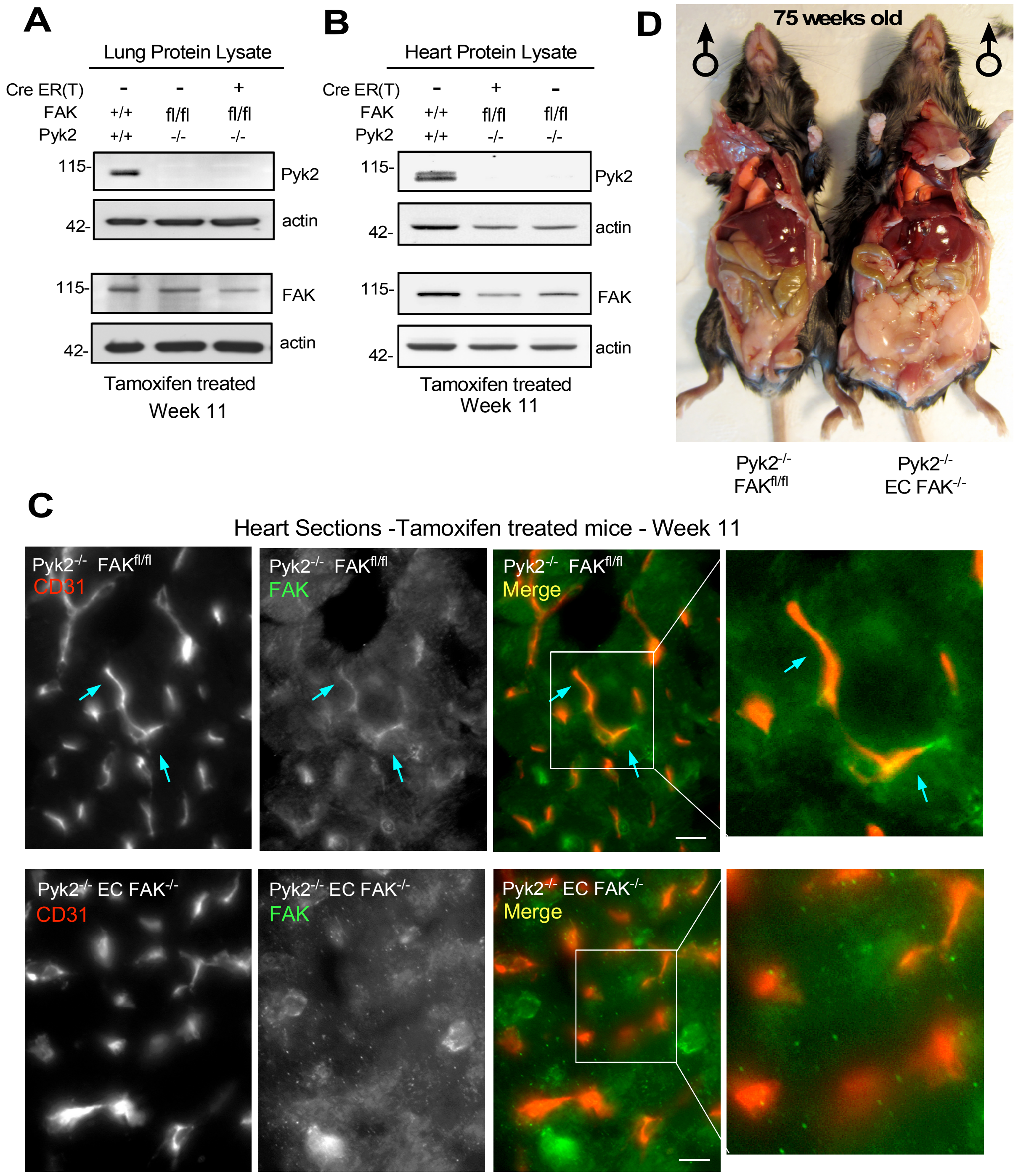


**Figure 1 - figure supplement 1.** Tamoxifen-induced EC FAK inactivation in Pyk2$^{-/-}$ FAK$^{fl/fl}$ Cre+ (Pyk2$^{-/-}$ EC FAK$^{-/-}$) mice. At 6 weeks of age (Cre- and Cre+) of the indicated genotypes were provided tamoxifen-chow for 3 weeks, rested one week on normal chow before analyses. (**A**) Lung or (**B**) heart protein lysates at ~11 weeks of age were analyzed by immunoblotting for Pyk2, FAK, and actin from mice with the indicated genotypes. (**C**) Frozen heart tissue sections from tamoxifen-treated Pyk2$^{-/-}$ FAK$^{fl/fl}$ and Pyk2$^{-/-}$ EC FAK$^{-/-}$ mice were analyzed by staining for ECs with CD31 (red) and FAK (green). Arrows (blue) indicate sites of EC-associated FAK staining overlap in Pyk2$^{-/-}$ FAK$^{fl/fl}$ mice. Scale is 20 µm. (**D**) Representative images of tamoxifen-treated Pyk2$^{-/-}$ FAK$^{fl/fl}$ and Pyk2$^{-/-}$ EC FAK$^{-/-}$ mice at 75 weeks of age.

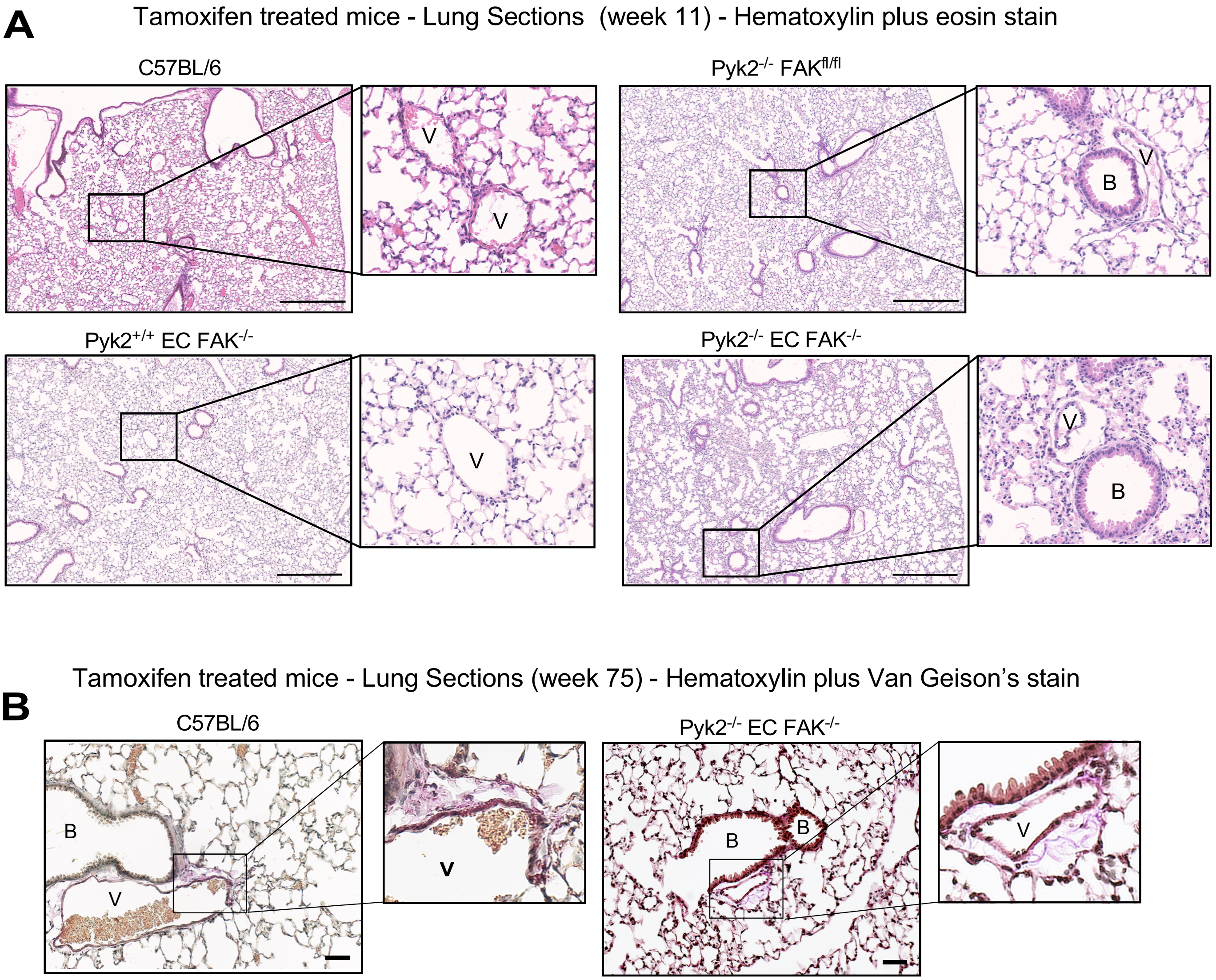


**Figure 1 - figure supplement 2.** Lung vessel changes upon EC FAK knockout in Pyk2$^{-/-}$ mice. (**A**) Representative H&E staining of formalin fixed and paraffin-embedded lung sections from tamoxifen-treated and saline perfused mice with the indicated genotypes. B, bronchiole and V, vessel. Scale is 500 μm. (**B**) Representative hematoxylin and Van Geison's lung section staining of 75 week old mice with the indicated genotypes. Some vessels contain red blood cells due to incomplete saline perfusion at euthanasia. Scale is 50 μm.

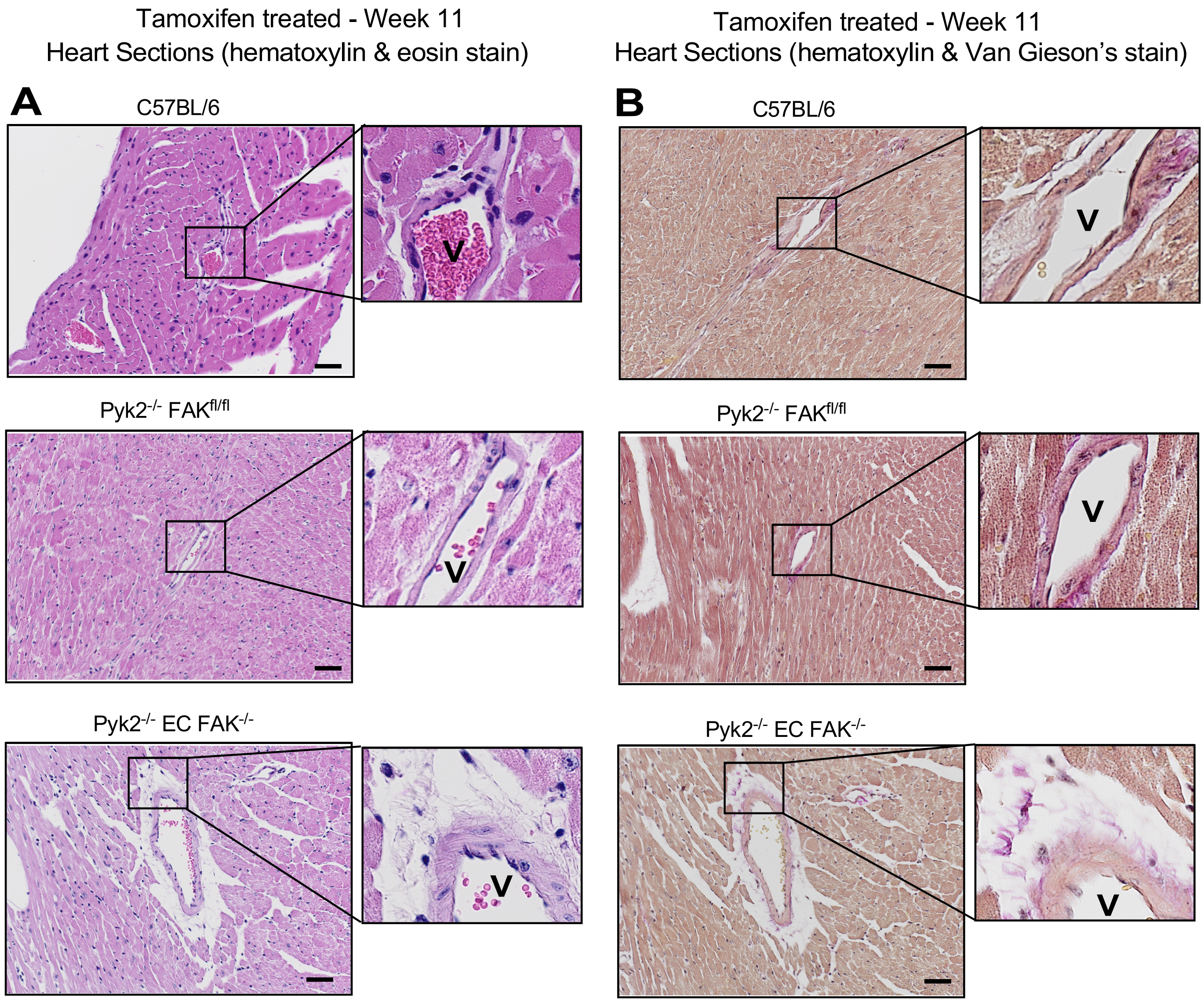


**Figure 1 - figure supplement 3.** Heart vessel changes upon induced EC FAK knockout in Pyk2-/- mice. (**A**) Representative H&E staining of formalin fixed and paraffin-embedded heart sections from tamoxifen-treated and perfused mice with the indicated genotypes at 11 weeks of age. B, bronchiole and V, vessel. (**B**) Representative hematoxylin and Van Geison's lung section staining of mice with the indicated genotypes at 11 weeks of age. Some vessels contain red blood cells due to incomplete saline perfusion at euthansia. Scale is 100 μm.

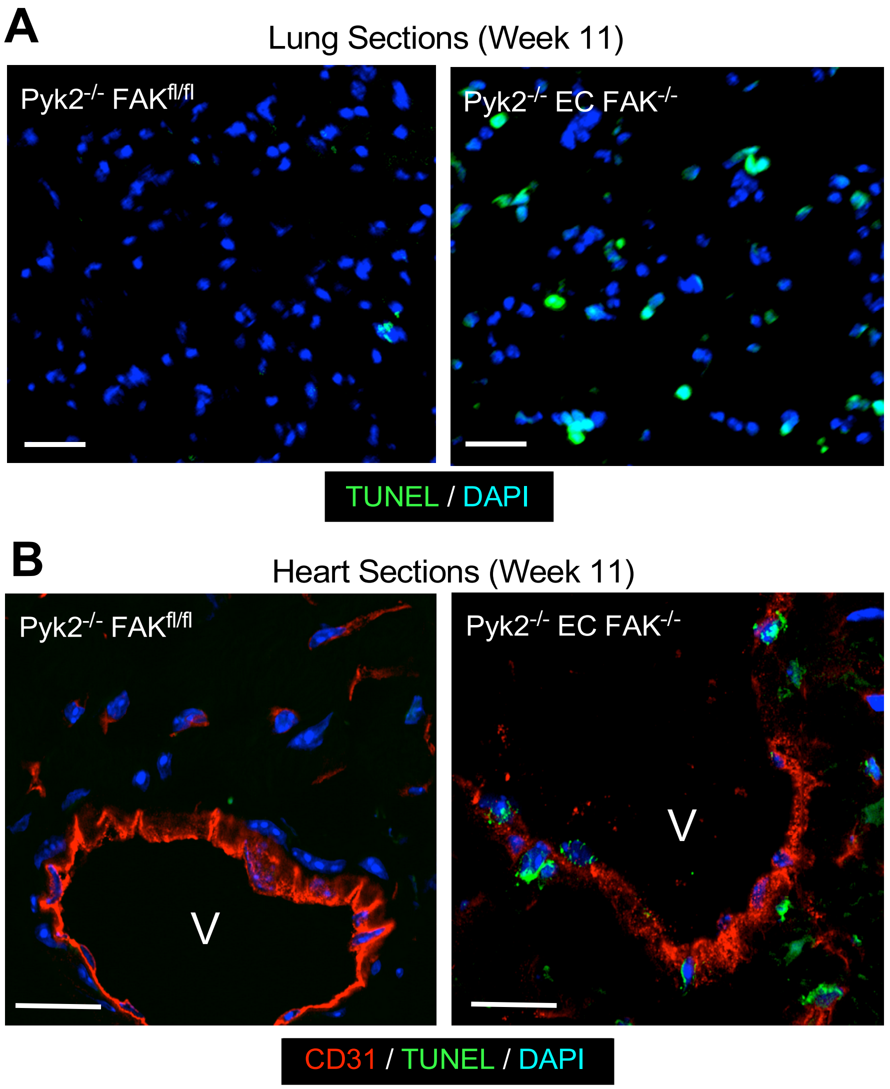


**Figure 3 - figure supplement 1.** Induced EC FAK knockout in Pyk2$^{-/-}$ mice triggers increased cell death *in vivo*. (**A**) Representative TUNEL (green) and DAPI (blue) immunostaining of lung frozen sections from tamoxifen-treated Pyk2$^{-/-}$ FAK$^{fl/fl}$ Cre- and Pyk2$^{-/-}$ EC FAK-/- (Cre+) mice. Scale is 20 µm. (**B**) Representative CD31 (red), TUNEL (green) and DAPI (blue) immunostaining of heart frozen sections from tamoxifen-treated Pyk2$^{-/-}$ FAK$^{fl/fl}$ and Pyk2$^{-/-}$ EC FAK$^{-/-}$ mice. Scale is 20 µm.

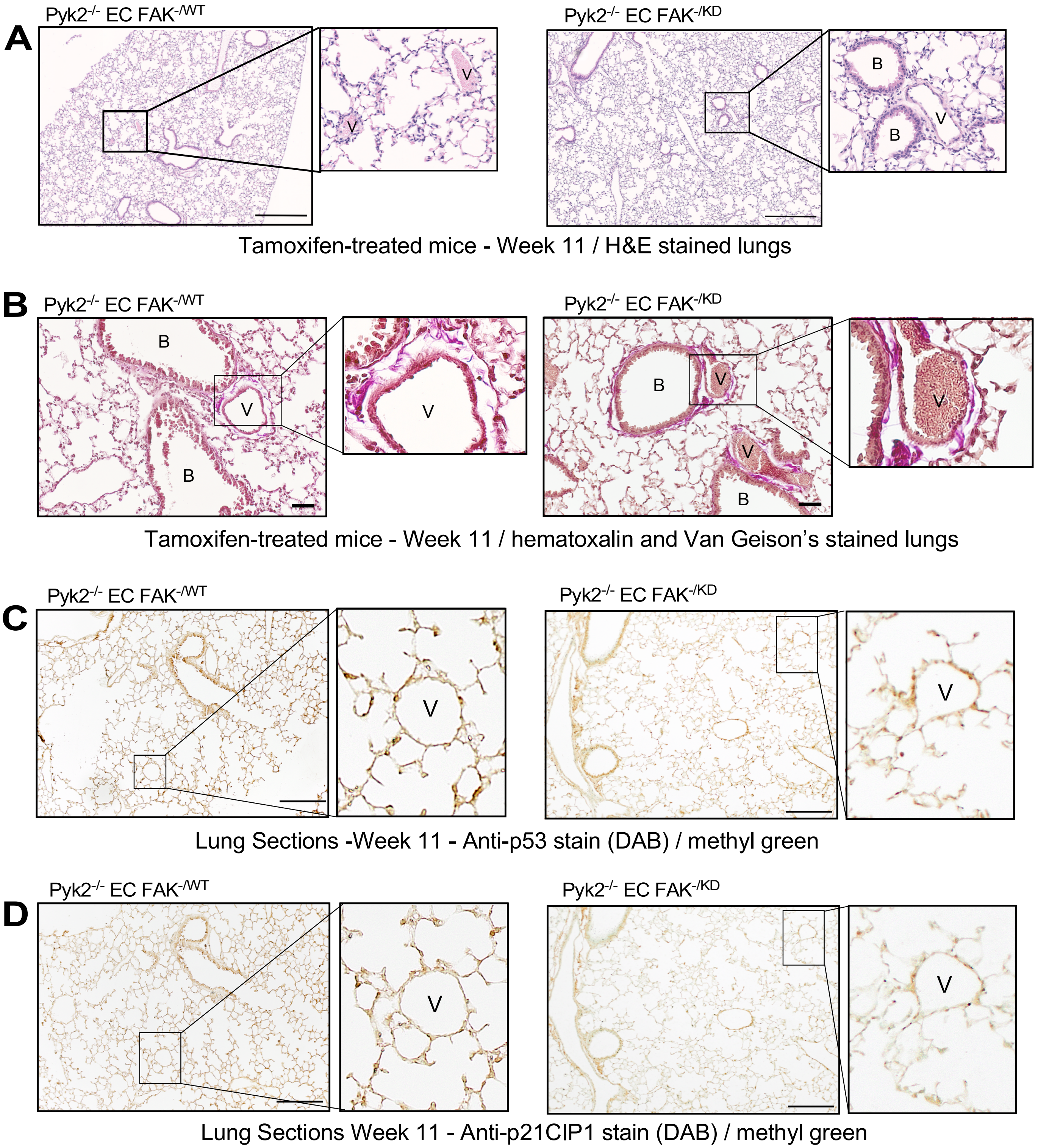


**Figure 4 - figure supplement 1.** No detectable differences in lung morphology, lung collagen deposition, or p53 and p21CIP1 staining in tamoxifen-treated Pyk2$^{-/-}$ EC FAK$^{-/WT}$ and Pyk2$^{-/-}$ EC FAK$^{-/KD}$ mice. (**A**) Representative H&E staining and (**B**) hematoxylin and Van Geison's staining of formalin fixed and paraffin-embedded lung sections. Scale is 200 µm. (**C**) Representative anti-p53 staining and (**D**) anti-p21CIP1 staining of paraffin-embedded lung sections of tamoxifen-treated mice with the indicated genotypes. Antibodies were detected by 3,3'-diaminobenzidine (DAB, brown) and slides were counter-stained with methyl green. Inset, lung blood vessel (V). Scale is 250 µm.

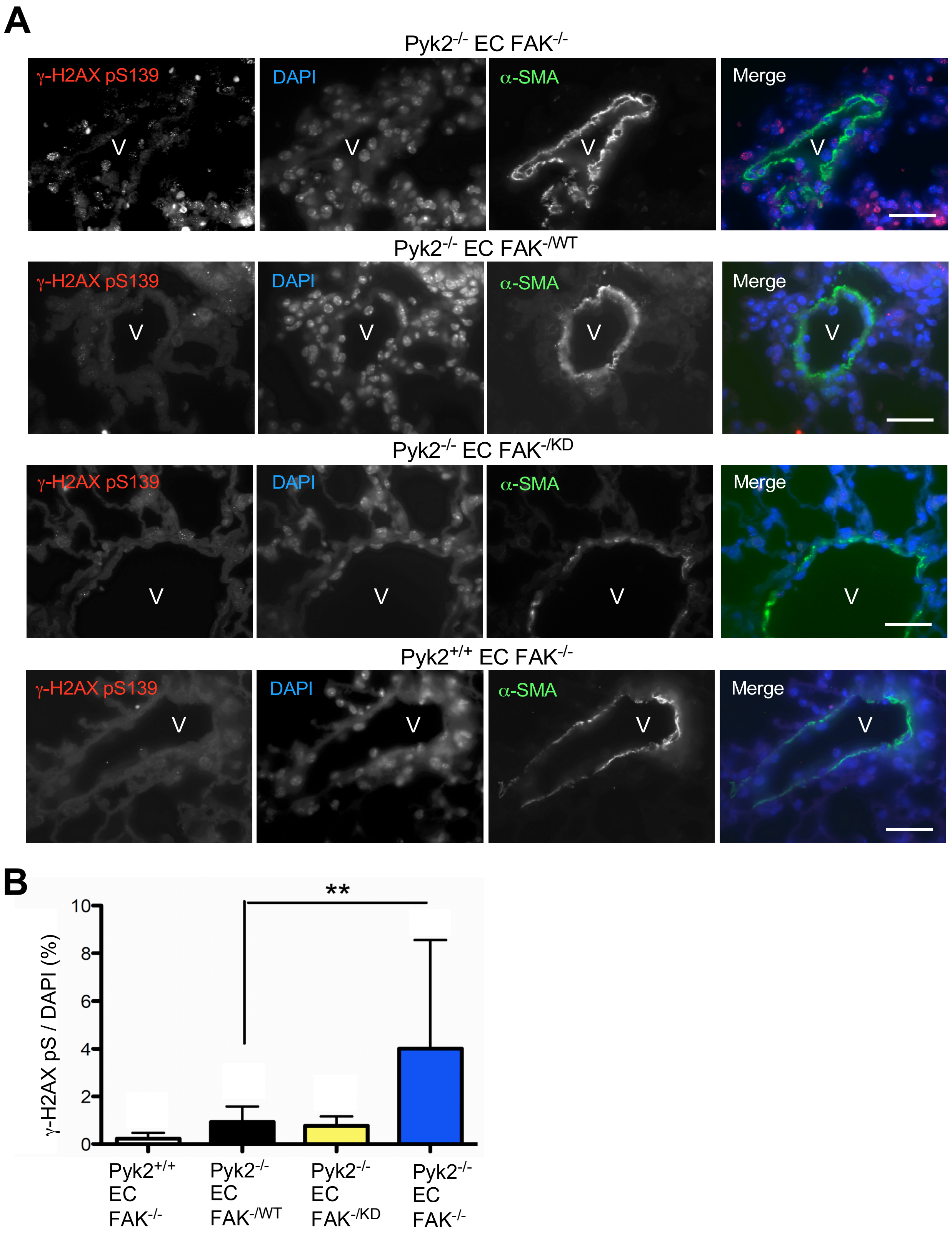


**Figure 4 - figure supplement 2.** Elevated DNA damage response marker staining upon EC FAK inactivation in Pyk2$^{-/-}$ lung tissue. (**A**) Representative γ-H2AX pS149 (red), nuclei (DAPI, blue), α-smooth muscle actin (α-SMA, green), and merged images of lung frozen sections from tamoxifen-treated Pyk2$^{+/+}$ EC FAK$^{-/-}$, Pyk2$^{-/-}$ EC FAK$^{-/-}$, Pyk2$^{-/-}$ EC FAK$^{-/WT}$, and Pyk2$^{-/-}$ EC FAK$^{-/KD}$ mice at week 11. Scale is 20 µm. (**B**) Quantification of immunostaining shown in panel A. Percentage of γ-H2AX pS149 and DAPI-double positive stained cells per field surrounding an α-SMA-positive vessel. Values are means from 10 images, n= 2 mice each genotype, +/- SD ** p<0.01, ANOVA with Tukey's post hoc test).

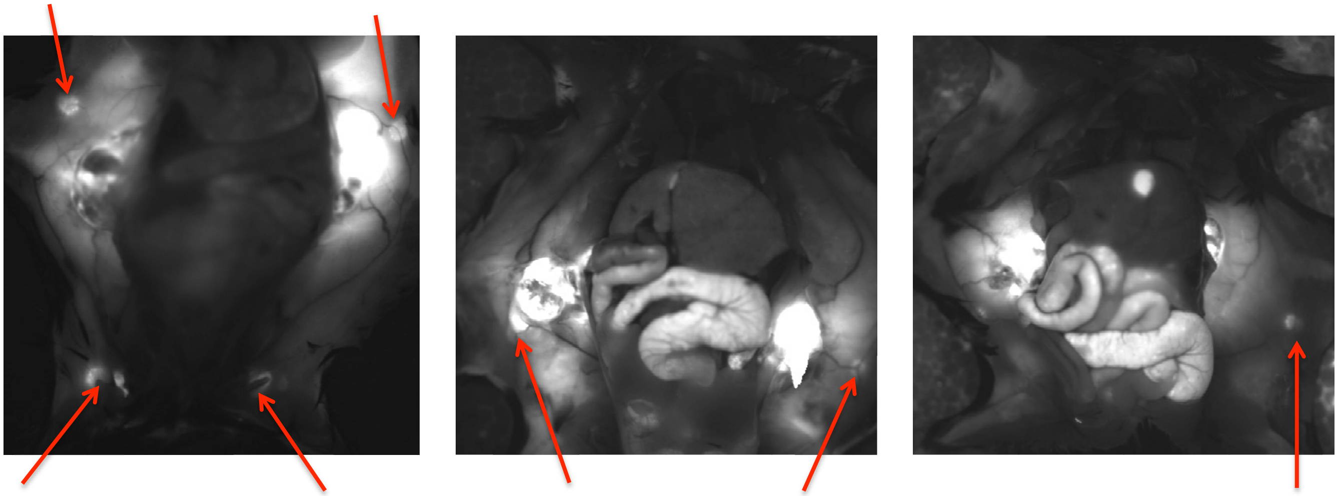

B16F10-RFP Lymph Node Metastasis

| | Axillary metastasis | Inguinal metastasis | total lymph nodes |
|---|---|---|---|
| C57BL/6 | 10/10 | 7/10 | 17/20 (85%) |

**Figure 5 - figure supplement 1.** Monitoring melanoma metastasis to lymph nodes. Detection of spontaneous B16F10-RFP metastasis at Day 21 to auxiliary and inguinal lymph nodes as detected by *in situ* fluorescent detection (Olympus, OV100). Values (percent of total) are numbers of tamoxifen-treated tumor-bearing mice C57Bl6 mice.

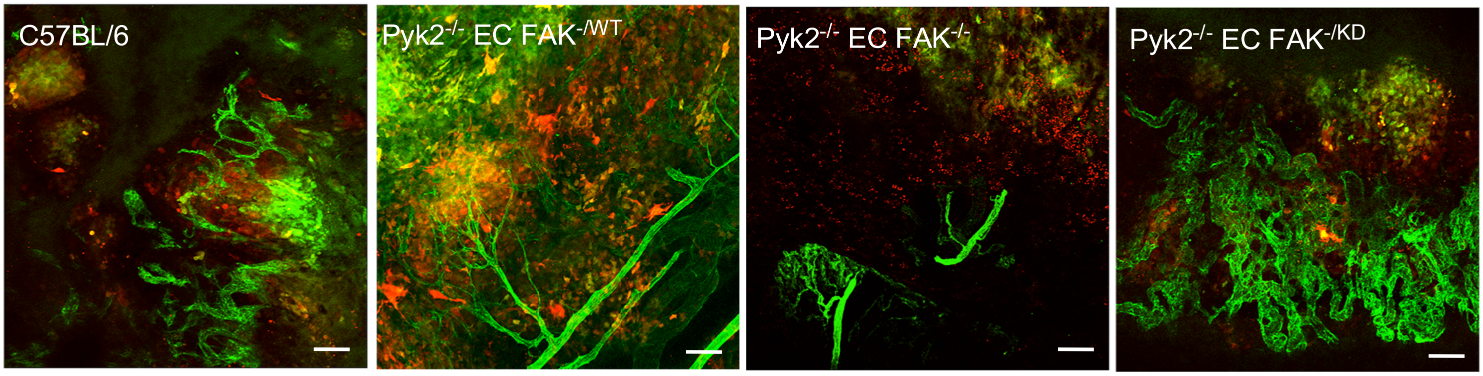


**Figure 5 - figure supplement 2.** Tumor-associated blood vessel organization requires EC FAK activity. RFP-labeled B16F10 melanoma cells in Matrigel were subcutaneously implanted in mice of the indicated tamoxifen treated mice for 7 days. Vasculature was labeled by tail vein FITC-lectin injection, Matrigel plugs were excised, and imaged by laser scanning confocal microscopy (Nikon). Shown are representative Z-stack images (10X) using 488 /568 channel series. Matrigel plugs from Pyk2$^{-/-}$ EC FAK$^{-/KD}$ mice exhibit disorganized vessels at plug periphery. Scale is 20 μm.

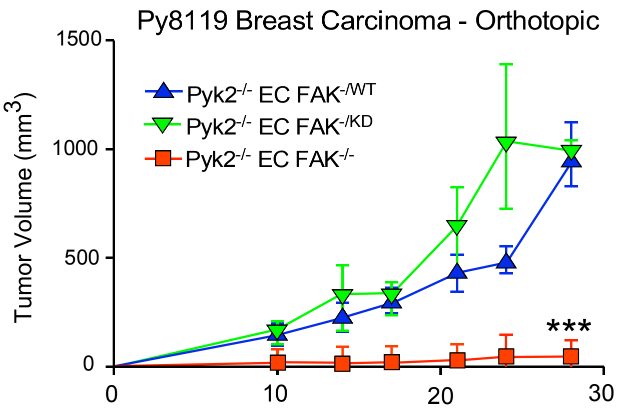


**Figure 5 - figure supplement 3.** Syngeneic Lewis lung carcinoma (LLC) tumor growth. 1 million LLC tumor cells were injected subcutaneous in tamoxifen-treated Pyk2$^{-/-}$ ECFAK$^{-/WT}$ (blue triangles), Pyk2$^{-/-}$ EC FAK$^{-/KD}$ (green triangles), and Pyk2$^{-/-}$ EC FAK$^{-/-}$ (red squares) mice. Values are tumor volume means by caliper measurements over 28 days. (n = 8 mice per group, +/- SD, *** P <0.001, ANOVA with Tukey's post hoc test).

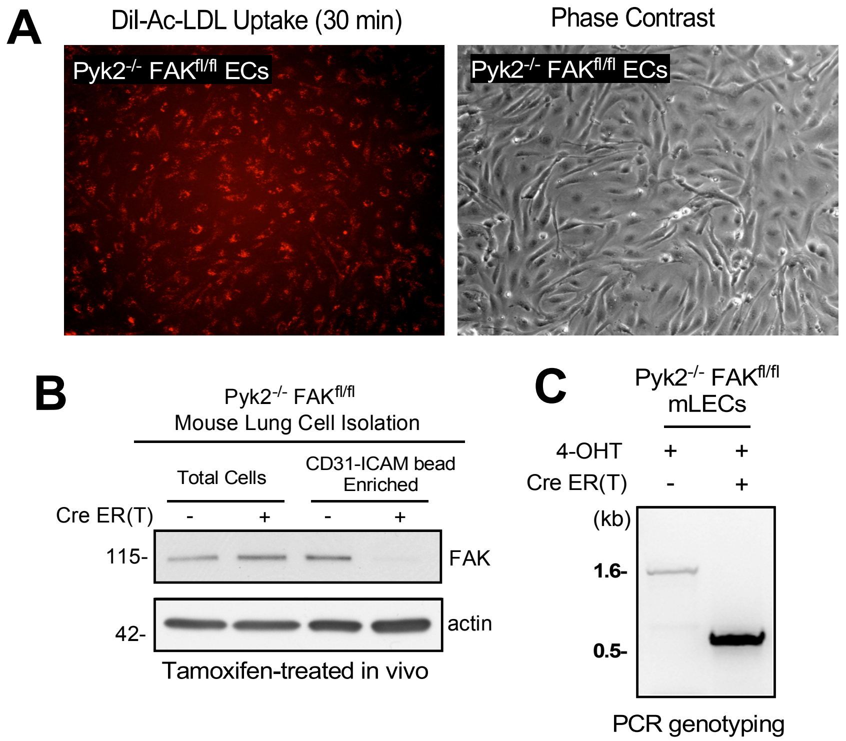


**Figure 7, figure supplement 1.** Primary mouse lung endothelial cells (MLECs). (**A**) Pyk2$^{-/-}$ FAK$^{fl/fl}$ MLECs were isolated by anti-CD31 and anti-ICAM2 affinity capture from 6-week-old mouse lung tissue and analyzed by acetylated low-density lipoprotein (DiI-Ac-LDL) uptake (red) and phase contrast (10X) microscopy. (**B**) Lysates of Pyk2$^{-/-}$ FAK$^{fl/fl}$ lung cells prior to (total cells) and after anti-CD31 / anti-ICAM2 affinity binding from tamoxifen-treated mice evaluated by immunoblotting for FAK and actin. (**C**) DNA genotyping from 4-hydroxy-tamoxifen (4-OHT) treated (48 h) Pyk2$^{-/-}$ FAK$^{fl/fl}$ Cre- or Cre+ MLECs. Cre-mediated recombination of floxed FAK yields a 500 bp product.

**Figure 4, Table supplement 1.** Phenotypes observed after tamoxifen treatment of SCL-Cre+ $FAK^{fl/WT}$ or $FAK^{fl/KD}$ mice on a $Pyk2^{-/-}$ C57BL/6 background at 11 weeks of age.

| Phenotype | EC $FAK^{-/WT}$ | EC $FAK^{-/KD}$ |
|---|---|---|
| Enlarged Lungs | 0/8 | 1/12 |
| Enlarged Heart | 0/8 | 0/12 |
| Enlarged Liver | 0/8 | 0/12 |
| Enlarged Kidneys | 0/8 | 0/12 |
| Death | 0/8 | 0/12 |
